# Synthetic Promoter Designs Enabled by a Comprehensive Analysis of Plant Core Promoters

**DOI:** 10.1101/2021.01.07.425784

**Authors:** Tobias Jores, Jackson Tonnies, Travis Wrightsman, Edward S. Buckler, Josh T. Cuperus, Stanley Fields, Christine Queitsch

## Abstract

Targeted engineering of plant gene expression holds great promise for ensuring food security and for producing biopharmaceuticals in plants. However, this engineering requires thorough knowledge of *cis*-regulatory elements in order to precisely control either endogenous or introduced genes. To generate this knowledge, we used a massively parallel reporter assay to measure the activity of nearly complete sets of promoters from Arabidopsis, maize and sorghum. We demonstrate that core promoter elements – notably the TATA-box – as well as promoter GC content and promoter-proximal transcription factor binding sites influence promoter strength. By performing the experiments in two assay systems, leaves of the dicot tobacco and protoplasts of the monocot maize, we detect species-specific differences in the contributions of GC content and transcription factors to promoter strength. Using these observations, we built computational models to predict promoter strength in both assay systems, allowing us to design highly active promoters comparable in activity to the viral 35S minimal promoter. Our results establish a promising experimental approach to optimize native promoter elements and generate synthetic ones with desirable features.

## Introduction

Precise control of gene expression is necessary to generate transgenic plants with new properties, such as growth in formerly incompatible environments or production of medically or nutritionally important products (Liu and Stewart, 2015; Lomonossoff and D’Aoust, 2016). Much of this control occurs at the initiation of transcription, the first committed step in gene expression. Transcription initiation involves the recruitment of the basal transcription machinery, comprised of general transcription factors and RNA polymerase, to core promoters. Core promoters define the transcription start site (TSS), but their activity typically leads to only low levels of expression (Smale and Kadonaga, 2003; Andersson and Sandelin, 2020). This basal level of transcription is increased by the interaction of core promoters with enhancers, which can reside upstream or downstream of the TSS and over a wide range of distances from the promoter (Banerji et al., 1981, 1983; Ricci et al., 2019).

The first core promoter element identified was the TATA-box. This motif, with the consensus sequence TATA(A/T)A(A/T), is recognized by the TATA-binding protein, a subunit of TFIID, and plays an important role in recruiting the basal transcription machinery and in determining the TSS location (Grosschedl and Birnstiel, 1980; Wasylyk et al., 1980; Smale and Kadonaga, 2003). Since then, several other core promoter elements have been discovered in viral and animal promoters (Grosschedl and Birnstiel, 1980; Smale and Baltimore, 1989; Ince and Scotto, 1995; Burke and Kadonaga, 1996; Lagrange et al., 1998; Lewis et al., 2000; Lim et al., 2004; Deng and Roberts, 2005; Parry et al., 2010). In plants, short motifs composed of pyrimidine bases, termed the TC motif or Y patch, have been described as potential plant-specific core promoter elements (Molina and Grotewold, 2005; Yamamoto et al., 2007; Bernard et al., 2010).

Apart from these elements, promoters also contain binding sites for transcription factors (TFs) close to the TSS. In contrast to the core promoter elements, which often occur at specific distances from and in a fixed orientation to the TSS, the TF binding sites can be functional in either orientation and their activity is less constrained by their distance to the TSS. Promoter-proximal TF binding sites can influence the transcriptional output from the nearby TSS and, in some cases, influence where transcription starts (Blake et al., 1990). In this study, we refer to the region surrounding the TSS that harbors core promoter elements as the core promoter; the extended region that includes the core promoter and upstream transcription factor binding sites is referred to as the promoter.

To gain a better understanding of the regulatory principles governing promoter activity, several high-throughput studies have been performed in yeast, *Drosophila melanogaster* and human cells (Patwardhan et al., 2009; Sharon et al., 2012; Lubliner et al., 2015; Arnold et al., 2017; van Arensbergen et al., 2017; Weingarten-Gabbay et al., 2019; de Boer et al., 2020; Kotopka and Smolke, 2020). These studies validated the contribution of core promoter elements and promoter-proximal TF binding sites to overall promoter activity and deduced rules governing the interaction among those elements. However, it is not clear whether these rules also apply to plant promoters. Although computational analyses have revealed that many of the core promoter elements identified in animals are enriched in plant promoters (Molina and Grotewold, 2005; Yamamoto et al., 2007; Kumari and Ware, 2013; Morton et al., 2014), only the TATA-box and the Initiator (Inr) element have been functionally validated (Zhu et al., 1995; Kiran et al., 2006; Srivastava et al., 2014; Jores et al., 2020). Some plant promoters do not harbor any of the known core promoter elements (Kumari and Ware, 2013). To date, large scale functional studies have not been performed with plant promoters.

A deeper understanding of the regulatory code of plant promoters and how it shapes transcription levels will further our knowledge of gene regulation, empower the controlled manipulation of gene expression for crop improvement and enable the rational design of promoters for use in genetic engineering. Here, we set out to comprehensively analyze the core promoters of the model plant Arabidopsis (*Arabidopsis thaliana*), and the important crop maize (*Zea mays*) and its close relative sorghum (*Sorghum bicolor*). The genome of the crucifer Arabidopsis is compact (∼135 Mb) and AT-rich, while the genomes of the cereals maize and sorghum are GC-rich and many times larger (∼2.7 Gb and ∼730 Mb, respectively). We sought to determine how these differences in genome content and architecture would be reflected in features of their promoter elements. Here, we identified key determinants of core promoter strength and characterized similarities and differences in the regulatory code of mono- and dicotyledonous plants. Using this knowledge, we designed synthetic core promoters with activities reaching levels comparable to that of the 35S minimal promoter. Furthermore, we trained computational models that accurately predict promoter strength in our assays and help improve promoter activity.

## Results

### Use of the STARR-seq assay to study plant core promoters

We used the self-transcribing active regulatory region sequencing (STARR-seq) assay, which we had established in plants (Jores et al., 2020), to measure the strength of nearly complete sets of core promoters from Arabidopsis, maize and sorghum. Specifically, for each species, we interrogated the sequences from −165 to +5 relative to the annotated TSS for protein-coding and microRNA (miRNA) genes. These 170 bp regions were tested for promoter strength by using them to drive expression of a barcoded green fluorescent protein (GFP) reporter gene (Fig. 1a). We included the first 5 bases after the TSS to cover core promoter elements that span the TSS, like the Inr, while avoiding substantial parts of the 5′ untranslated region (UTR). 5′ UTRs affect mRNA levels post-transcriptionally and hence their inclusion could confound assessment of promoter strength (Srivastava et al., 2018). Instead, we used the 5′ UTR of a sorghum histone H3 gene (SORBI_3010G047100) for all sorghum promoters and the 5′ UTR of a maize histone H3.2 gene (Zm00001d041672) for all maize and Arabidopsis promoters (the 5′ UTR of the Arabidopsis histone H3.1 gene AT5G10390 had intrinsic promoter activity). We constructed three STARR-seq libraries that contained 18,329 Arabidopsis, 34,415 maize and 27,094 sorghum core promoters linked to approximately 400,000 unique barcodes per library (Supplementary Table 1). To test these promoters for their response to a strong enhancer, we also generated each library using a plasmid containing the cauliflower mosaic virus 35S enhancer (Fang et al., 1989; Benfey et al., 1990) immediately upstream of the promoter insertion site (Jores et al., 2020). The six libraries were assayed individually in transiently transformed tobacco leaves and maize protoplasts.

**Fig. 1.**
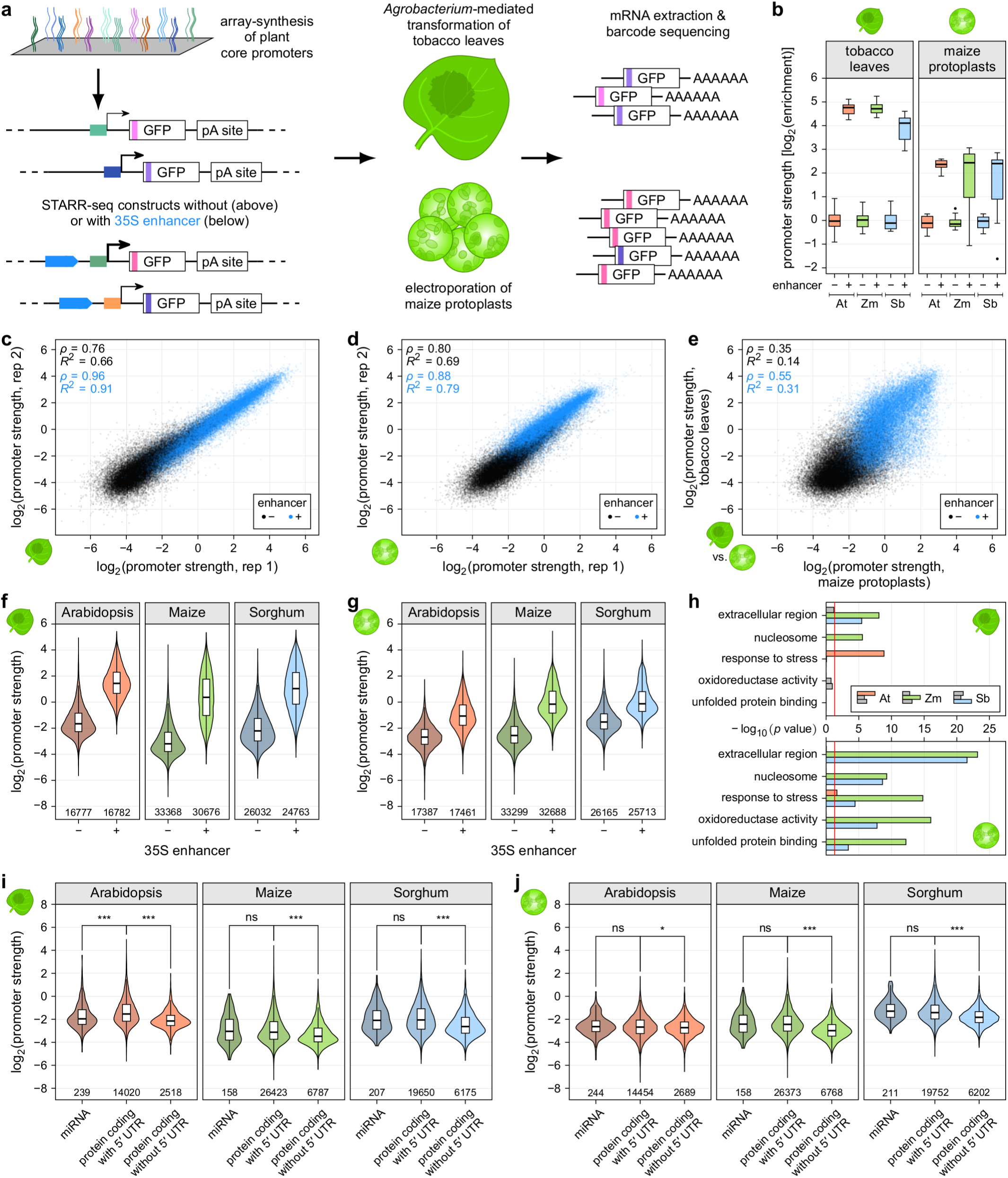
STARR-seq measures core promoter strength in tobacco leaves and maize protoplasts. **a**, Assay scheme. The core promoters (bases −165 to +5 relative to the TSS) of all genes of Arabidopsis, maize and sorghum were array-synthesized and cloned into STARR-seq constructs to drive the expression of a barcoded GFP reporter gene. For each species, two libraries, one without and one with a 35S enhancer upstream of the promoter, were created. The libraries were subjected to STARR-seq in transiently transformed tobacco leaves and maize protoplasts. **b**, Each promoter library (At, Arabiopsis; Zm, maize; Sb, sorghum) contained two internal control constructs driven by the 35S minimal promoter without (−) or with (+) an upstream 35S enhancer. The enrichment (log_2_) of recovered mRNA barcodes compared to DNA input was calculated with the enrichment of the enhancer-less control set to 0. In all following figures this metric is indicated as promoter strength. Each boxplot (center line, median; box limits, upper and lower quartiles; whiskers, 1.5 × interquartile range; points, outliers) represents the enrichment of all barcodes linked to the corresponding internal control construct. **c**,**d**, Correlation (Pearson’s *R*^2^ and Spearman’s *ρ*) of two biological replicates of STARR-seq using the maize promoter libraries in tobacco leaves (**c**) or in maize protoplasts (**d**). **e**, Comparison of the strength of maize promoters in tobacco leaves and maize protoplasts. Pearson’s *R*^2^ and Spearman’s *ρ* are indicated. **f**,**g**, Violin plots of the strength of plant promoters from the indicated species as measured by STARR-seq in tobacco leaves (**f**) or maize protoplasts (**g**) for libraries without (−) or with (+) the 35S enhancer upstream of the promoter. **h**, Enrichment of selected GO terms for genes associated with the 1000 strongest promoters in the Arabidopsis (At), maize (Zm), and sorghum (Sb) promoter libraries without enhancer in tobacco leaves (top panel) and maize protoplasts (bottom panel). The red line marks the significance threshold (adjusted *p* value ≤ 0.05). Non-significant bars are shown in gray. **i**,**j**, Violinplots of promoter strength (libraries without 35S enhancer) in tobacco leaves (**i**) or maize protoplasts (**j**). Promoters were grouped by gene type. In all figures, violinplots represent the kernel density distribution and the boxplots within represent the median (center line), upper and lower quartiles (box limits), and 1.5 × the interquartile range (whiskers) for all corresponding promoters. Numbers at the bottom of the plot indicate the number of tested promoters. Significant differences between two samples were determined using the Wilcoxon rank-sum test and are indicated: ∗, *p* ≤ 0.01; ∗∗, *p* ≤ 0.001; ∗∗∗, *p* ≤ 0.0001; ns, not significant.

In each promoter library, we included two control constructs, one containing only the viral 35S minimal promoter (-46 to +5 relative to the TSS) and the other containing the 35S minimal promoter and enhancer (-199 to -47 relative to the TSS). The promoter strength for each tested plant promoter was normalized to the control construct containing only the 35S minimal promoter. The construct also containing the strong 35S enhancer upstream of the minimal promoter was used to test the dynamic range of the assay. Consistent with previous reports (Bruce et al., 1989; Jores et al., 2020), the 35S enhancer was four-fold more active in the tobacco system than in maize protoplasts (Fig. 1b). We performed two biological replicates for each promoter library in each assay system. The replicates were highly correlated, especially for the libraries with the 35S enhancer, which reflected their generally higher promoter strength (Fig. 1c,d and Supplementary Fig. 1). Therefore, we used the average promoter strength from both replicates for all further analyses. We validated these results by re-testing a subset of 166 and 173 promoters in two separate libraries, obtaining results that were highly correlated with the data from the comprehensive promoter libraries (Supplementary Fig. 2). Since the sorghum promoters were coupled to a sorghum 5′ UTR in the comprehensive library and to a maize 5′ UTR in the validation libraries, the high correlation between these datasets suggests that the two 5′ UTRs did not strongly affect promoter strength.

Promoter strengths as measured in the tobacco leaf system had a weak to intermediate (*R*^2^ of 0.14 to 0.40) correlation with those obtained from maize protoplasts (Fig. 1e and Supplementary Fig. 1c,f), indicating that there are substantial differences in how the two systems interact with the core promoters. Irrespective of the assay system, the promoters spanned a wide range of activity, with over 250-fold difference between the strongest and weakest promoters (Fig. 1f,g and Supplementary Table 2). Few promoters were stronger than the viral 35S minimal promoter, which is likely optimized for maximal activity. Overall, the promoters of the dicot Arabidopsis tended to perform better in the dicot tobacco system, while the promoters of the monocots maize and sorghum showed greater activity in protoplasts of the monocot maize (Fig. 1f,g).

GO term enrichment analysis showed that the genes corresponding to the most active promoters in our assay were significantly (adjusted *p* value ≤ 0.05) enriched for components of nucleosomes, which are highly expressed housekeeping genes (Fig. 1h). In both systems, strong promoters often were also associated with genes annotated for response to stress and function in the extracellular region, including genes encoding defense and cell wall proteins. In the maize protoplast system, genes associated with strong promoters frequently encoded proteins with oxidoreductase activity or unfolded protein binding functions. The latter is consistent with reports of wound-induced reactive oxygen species and a heat shock response in protoplasts (Yahraus et al., 1995). Although these results show a qualitative agreement between core promoter strength and expression level for some genes, there was no significant correlation overall between promoter strength and expression data (Walley et al., 2016; Wang et al., 2018; Mergner et al., 2020) for the corresponding genes *in planta* (Supplementary Fig. 3). This lack of correlation is expected, as core promoters represent only a subset of all the regulatory elements that drive gene expression, and other elements such as enhancers can drastically affect transcription rates in the genomic context.

Next, we asked if genes of different types employ different promoters. The activity of miRNA promoters was indistinguishable from that of promoters of protein-coding genes (Fig. 1i,j). However, promoters from genes with an annotated 5′ UTR were generally stronger than those of genes without a 5′ UTR annotation. As the TSSs of the latter are probably not correctly annotated, these sequences are likely not true promoters, explaining their low activity.

### GC content, core promoter elements and transcription factor binding sites influence promoter strength

Monocot genomes are more GC-rich than dicot genomes (Kumari and Ware, 2013; Singh et al., 2016), and this bias holds true for their core promoter sequences (Fig. 2a). In the tobacco leaf system, GC content strongly affected promoter strength, with AT-rich promoters up to four-fold more active than GC-rich ones (Fig. 2b). A high GC content was especially detrimental close to the 5′ end of the promoters but was better tolerated towards the 3′ end (Fig. 2c). In contrast, in maize protoplasts, GC content was not predictive of promoter strength (Fig. 2d). Since the GC content of the Arabidopsis and tobacco genomes is similar (Rensink et al., 2005), the transcriptional machinery in tobacco is likely tuned to AT-rich promoters and works less well with the GC-rich promoters of maize and sorghum. Conversely, the transcription machinery of maize commonly acts on GC-rich promoters and can effectively use them in protoplasts. The correlation between promoter strength and GC content is, therefore, a characteristic of the assay system and not an intrinsic feature of the promoters.

**Fig. 2.**
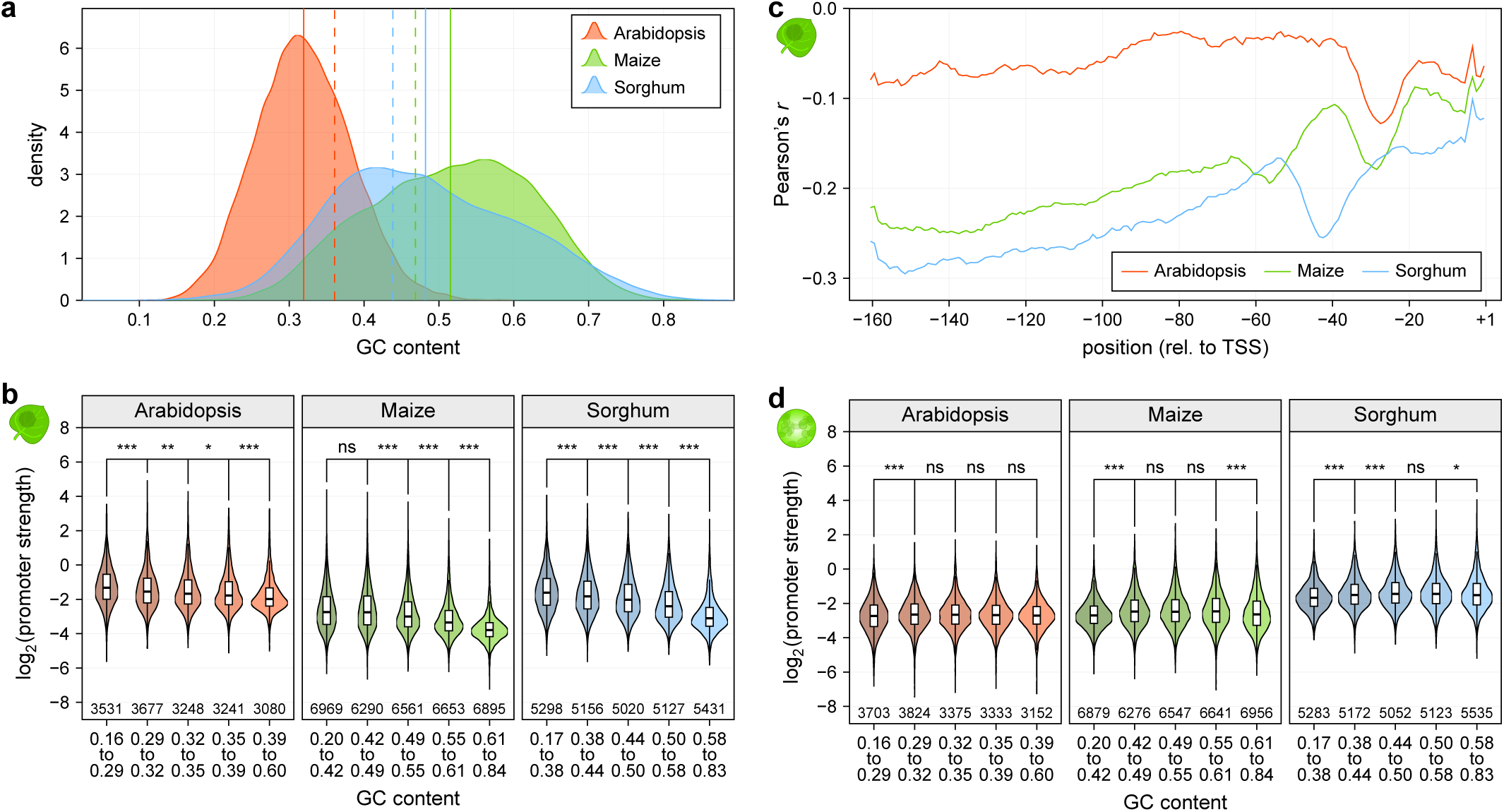
GC content affects promoter strength in tobacco leaves. **a**, Distribution of GC content for all promoters of the indicated species. Lines denote the mean GC content of promoters (solid line) and the whole genome (dashed line). **b**, Violin plots (as defined in Figure 1) of promoter strength for libraries without enhancer in tobacco leaves. Promoters are grouped by GC content to yield groups of approximately similar size. **c**, Correlation (Pearson’s *r*) between promoter strength and the GC content of a 10 base window around the indicated position in the plant promoters. **d**, As (**b**) but for promoter strength in maize protoplasts.

We next tested how known core promoter elements affect promoter strength. Considering first the location of TATA-box motifs, we noticed marked differences among the promoters of Arabidopsis, maize and sorghum. In Arabidopsis promoters, the distribution of TATA-boxes had a peak ∼30 bp upstream of the TSS (Fig. 3a). Although this location also is common for maize promoters, the maize promoters showed two additional peaks for the TATA-box at: ∼55 and ∼70 bp upstream of the TSS. In sorghum promoters, the TATA-box distribution peaked at ∼40 bp upstream of the TSS, with a shoulder ∼30 bp upstream of the TSS.

**Fig. 3.**
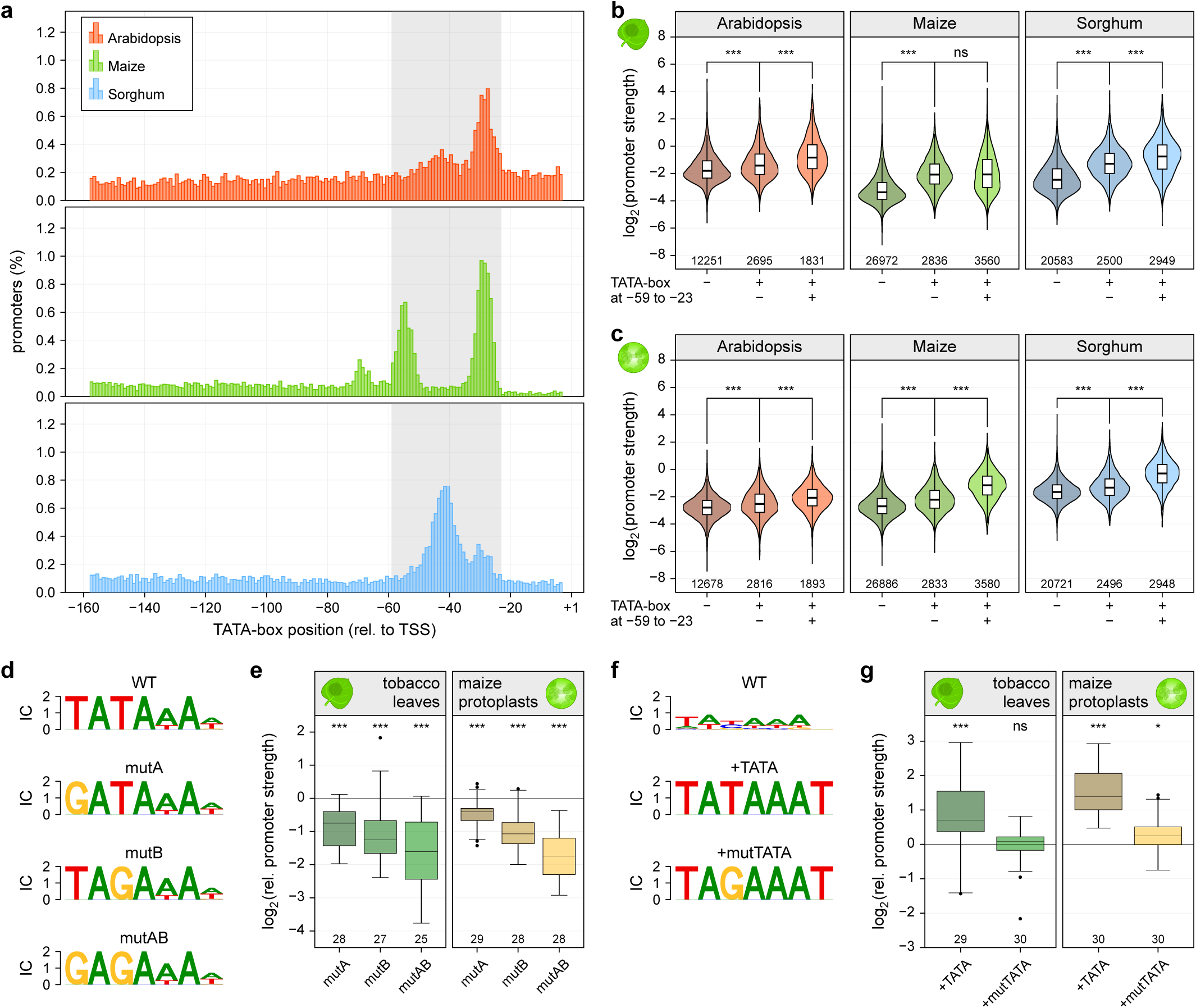
The TATA-box is a key determinant of promoter strength. **a**, Histograms showing the percentage of promoters with a TATA-box at the indicated position. The region between positions −59 and −23 in which most TATA-boxes reside is highlighted in gray. **b**,**c**, Violin plots (as defined in Figure 1) of promoter strength for libraries without enhancer in tobacco leaves (**b**) or maize protoplasts (**c**). Promoters without a TATA-box (−) were compared to those with a TATA-box outside (+/−) or within (+/+) the −59 to −23 region. **d**-**g**, Thirty plant promoters with a strong (**d,e**) or weak (**f,g**) TATA-box (WT) were tested. One (mutA and mutB) or two (mutAB) T>G mutations were inserted into promoters with a strong TATA-box (**d,e**). A canonical TATA-box (+TATA) or one with a T>G mutation (+mutTATA) was used to replace the weak TATA-box (**f,g**). Logoplots (**f,d**) of the TATA-box regions of these promoters and their strength (**g,e**) relative to the WT promoter (set to 0, horizontal black line) are shown. Boxplots (center line, median; box limits, upper and lower quartiles; whiskers, 1.5 × interquartile range; points, outliers) denote the strength of the indicated promoter variants. Numbers at the bottom of the plot indicate the number of tested promoter elements. Significant differences from a null distribution were determined using the Wilcoxon rank-sum test and are indicated: ∗, *p* ≤ 0.01; ∗∗, *p* ≤ 0.001; ∗∗∗, *p* ≤ 0.0001; ns, not significant.

Core promoters harboring a TATA-box were up to four-fold stronger than TATA-less ones, especially when the TATA-box is located within the region from 23 to 59 bp upstream of the TSS, where most TATA-boxes in the promoters of Arabidopsis, maize and sorghum reside (Fig. 3a-c). The location of the TATA-box in maize promoters affected their strength only in maize protoplasts. In this assay system, maize promoters with a TATA-box in one of the three peaks of the TATA-box distribution were stronger than those with a TATA-box elsewhere. Furthermore, maize promoters with a TATA-box in the peak closest to the TSS were strongest, and they became successively weaker in the other two peaks as the TATA-box is located increasingly more TSS-distal (Supplementary Fig. 4). The effect of the TATA-box on promoter strength was not a consequence of an increased AT-content in the promoters containing a TATA-box. (Supplementary Fig. 5). To directly measure the effect of the TATA-box, we mutated this motif in native promoters. Replacement of one or both T nucleotides in the core TATA motif with a G resulted in decreased transcriptional activity (Fig. 3d, e). Similarly, promoter strength was increased when a canonical TATA-box was inserted into a TATA-less promoter; a mutated version of the TATA-box did not have this effect (Fig. 3f, g).

In animal promoters, the TATA-box is often surrounded by the upstream (BRE^u^) and/or downstream (BRE^d^) TFIIB recognition element. Mutational studies have demonstrated that these elements can modulate promoter strength (Lagrange et al., 1998; Deng and Roberts, 2005). In tobacco leaves, neither of the two elements had a strong effect on promoter activity; however, in maize protoplasts, BRE^u^ was associated with 25% increased and BRE^d^ with 10% decreased promoter strength (Supplementary Fig. 6a-d and Supplementary Table 4). Consistent with these results, mutations that inactivate BRE^u^ decreased promoter strength in maize protoplasts but not in tobacco leaves. Inserting a canonical BRE^u^ led to increased promoter activity, especially in maize protoplasts. In contrast, mutating or inserting BRE^d^ had only modest effects on promoter activity in both assay systems (Supplementary Fig. 6e-h). A valine residue in the helix-turn-helix motif of the general transcription factor TFIIB is crucial for the recognition of BRE^u^ in animals (Lagrange et al., 1998; Tsai and Sigler, 2000). Although this residue is not conserved in any plant TFIIB protein, the maize genome encodes an additional TFIIB-related protein with a valine at the corresponding position (Supplementary Fig. 7). The presence of this maize-specific TFIIB-related protein may explain the increased activity of BRE^u^ in the maize protoplast system.

Computational analyses of plant promoters (Molina and Grotewold, 2005; Yamamoto et al., 2007; Bernard et al., 2010) have detected an enrichment of short, pyrimidine-rich motifs upstream of the TSS (Supplementary Fig. 8a). Because such an enrichment was not detected in animal promoters, these motifs, termed Y patches, were proposed to be plant-specific core promoter elements. Our data support this hypothesis, as Y patch-containing promoters showed 10 to 15% greater strength compared to those without the element (Supplementary Fig. 8b,c and Supplementary Table 4).

Consistent with previous studies (Zhu et al., 1995; Srivastava et al., 2014), we observed that promoters with an Inr at the TSS were generally stronger than those without it. In contrast, the polypyrimidine initiator TCT, previously described in animals (Parry et al., 2010), was less effective (Supplementary Fig. 9).

Finally, we asked whether promoter-proximal TF binding sites affect promoter strength. We first clustered TFs by similarity of their binding site motifs and created a consensus motif for each of the 72 clusters (Supplementary Table 3). We then compared the strength of promoters with a predicted binding site to that of promoters lacking it. About 67% of the TF clusters did not have a significant impact on promoter strength. However, 23 TF motifs were significantly (*p* value ≤ 0.0005) associated with altered promoter strength in at least one assay system (Supplementary Table 4). For example, the TCP transcription factor motif tends to reside in promoters that were strong in tobacco leaves, while this effect was not observed in maize protoplasts (Supplementary Fig. 10a,b). On the other hand, promoters with a motif for heat-shock factors (HSFs) were stronger than those without it in maize protoplasts but not in tobacco leaves (Supplementary Fig. 10c,d).

We asked whether core promoter elements and TF binding sites are spatially constrained in relation to one another. In contrast to core promoter elements, most TF binding sites did not show a preferential position relative to the TSS. However, we observed that TF binding sites upstream of the TATA-box were generally associated with a higher promoter strength compared to those downstream of the TATA-box (Supplementary Fig. 11 and Supplementary Table 5). Since RNA polymerase is recruited to the region downstream of the TATA-box, this enzyme may displace TFs bound here and thereby prevent them from activating transcription.

### Core promoters show varying degrees of enhancer responsiveness

In animals, promoters can interact differentially with enhancers (Gehrig et al., 2009; Arnold et al., 2017). Similarly, the 35S enhancer activated some plant core promoters more than others. However, the presence of the 35S enhancer resulted in increased transcription from almost all core promoters, up to 60-fold for the most responsive promoters in the tobacco leaf system and up to 15-fold in maize protoplasts; the 35S enhancer is less active in maize protoplasts (Bruce et al., 1989; Jores et al., 2020). Consistent with the notion that enhancers are the drivers of tissue- and condition-specific transcription (Benfey et al., 1990; Andersson and Sandelin, 2020), promoters of genes with high tissue-specificity (top third of the genes as ranked by the tissue-specificity index τ (Yanai et al., 2005)) showed on average 33% increased enhancer responsiveness compared to promoters of genes with low tissue-specificity (bottom third of the τ distribution) (Fig. 4a,b). Similarly, promoters of miRNA genes, which are often differentially expressed in response to environmental or developmental cues, were 33% more responsive to the 35S enhancer than promoters of protein-coding genes (Supplementary Fig. 12).

**Fig. 4.**
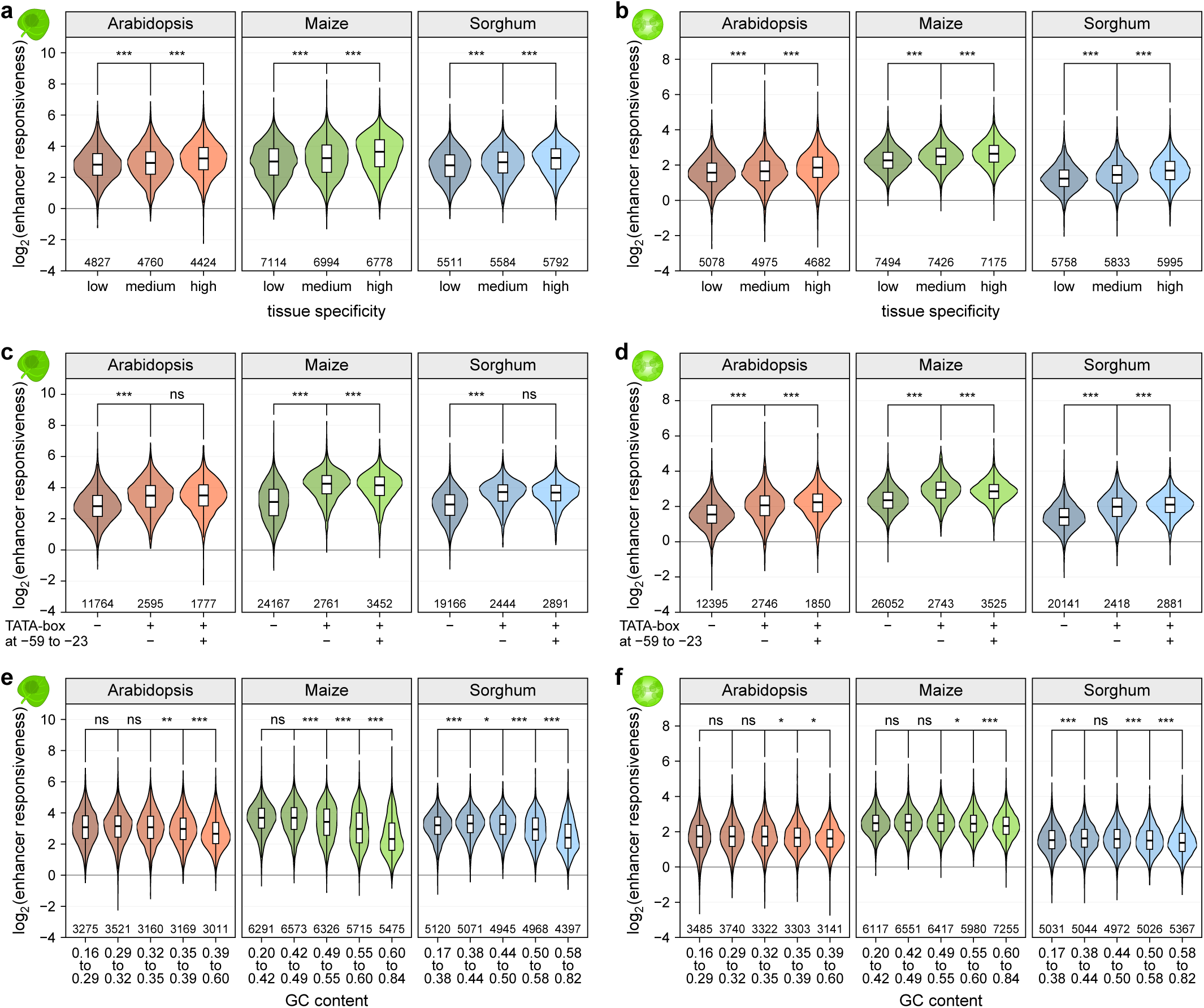
Enhancer responsiveness of promoters depends on the TATA-box and GC content. **a**,**b**, Violin plots (as defined in Figure 1) of enhancer responsiveness (promoter strength^with enhancer^ divided by promoter strength^without enhancer^) in tobacco leaves (**a**) or maize protoplasts (**b**). Promoters were grouped into three bins of approximately similar size according to the tissue-specificity *τ* of the expression of the associated gene. **c**,**d**, Violin plots of enhancer responsiveness in tobacco leaves (**c**) or maize protoplasts (**d**). Promoters without a TATA-box (−) were compared to those with a TATA-box outside (+/−) or within (+/+) the −59 to −23 region. **e**,**f**, Violin plots of enhancer responsiveness in tobacco leaves (**e**) or maize protoplasts (**f**) for promoters grouped by GC content.

To understand which promoter features influence enhancer responsiveness, we analyzed the elements that affect promoter strength. Promoters with a TATA-box were up to 67% more responsive to the 35S enhancer than TATA-less promoters; however, the location of the TATA-box did not have a consistent impact on enhancer responsiveness (Fig. 4c,d). Furthermore, promoter GC content influenced enhancer responsiveness in the tobacco leaf system, but not in maize protoplasts (Fig. 4e,f). While the GC content and TATA-box had a similar effect on enhancer responsiveness as on promoter strength, the same was not true for TFs. Instead, TFs that increased promoter strength often reduced enhancer responsiveness (Supplementary Fig. 13a-d), potentially due to competition for a limited pool of TFs or because of incompatibilities between recruited downstream factors. In contrast, some TFs that did not influence promoter strength affected enhancer responsiveness (Supplementary Fig. 13e,f). The effects on enhancer responsiveness possibly reflect synergistic effects, whereby the core transcriptional machinery and the TFs at promoters and enhancers interact with one another.

### Core promoter strength can be modulated by light

The plant STARR-seq assay can identify light-responsive enhancers (Jores et al., 2020). To test whether core promoters that respond to light can also be identified, we subjected the promoter libraries to STARR-seq experiments in tobacco leaves that were kept in the light (16h light, 8h dark) for two days after transformation (Fig. 5a). We did not perform the same experiment with maize protoplasts, as known light-responsive enhancers were not active in this system (Supplementary Fig. 14). As expected, most promoters did not respond to the light. However, about 2400 promoters were at least four times more active in the light or in the dark (Fig. 5b). The genes associated with the most highly light-dependent promoters were enriched for those encoding plastid proteins, especially for proteins in thylakoids, the membrane-bound chloroplast compartments that are the site of the light-dependent reactions of photosynthesis (Fig. 5c).

**Fig. 5.**
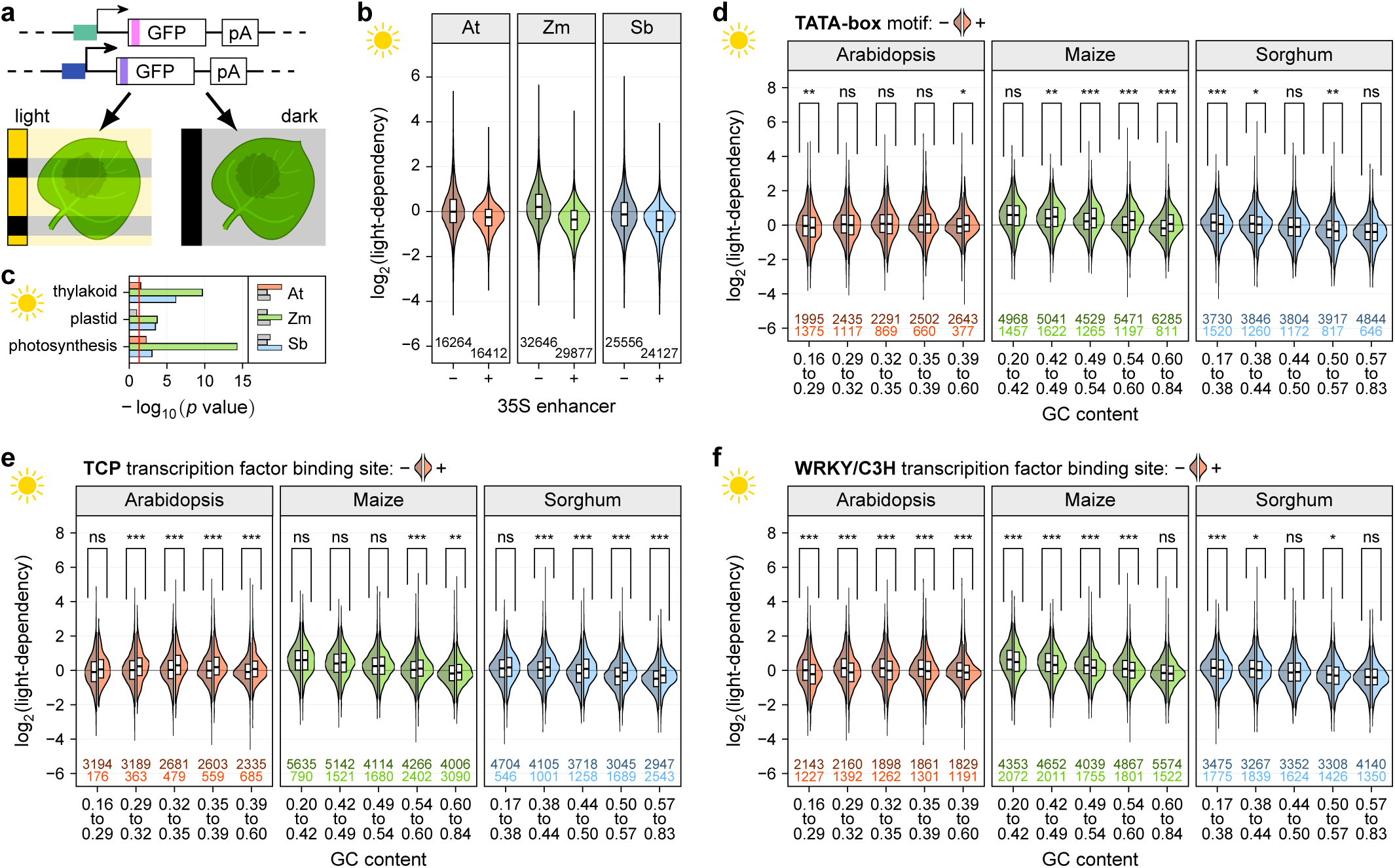
Promoter strength can be modulated by light. **a**, Tobacco leaves were transiently transformed with STARR-seq promoter libraries and the plants were kept for two days in 16h light/8h dark cycles (light) or completely in the dark (dark) prior to mRNA extraction. **b**, Violin plots (as defined in Figure 1) of light-dependency (promoter strength^light^ divided by promoter strength^dark^) for promoters in the libraries with (+) or without (−) the 35S enhancer. **c**, Enrichment of selected GO terms for genes associated with the 1000 most light-dependent promoters. The red line marks the significance threshold (adjusted *p* value ≤ 0.05). Non-significant bars are gray. **d**-**f**, Violin plots of light-dependency. Promoters are grouped by GC content and split into promoters without (left half, darker color) or with (right half, lighter color) a TATA-box (**d**), or a binding site for TCP (**e**) or WRKY (**f**) transcription factors.

While promoters that are AT-rich were more light-dependent than GC-rich ones (Fig. 5d), the effects of GC content on light-dependency were much less pronounced than on promoter strength and enhancer responsiveness. Similarly, the presence of a TATA-box showed weaker and even inconsistent effects on light-dependency compared to TATA-box effects on promoter strength and enhancer responsiveness (Fig. 5d). We found that the light-dependency of a promoter was mainly determined by the TF binding sites it contains. The presence of the TCP binding site, for example, led to increased expression in the light (Fig. 5e) and, consistent with previous studies (Heerah et al., 2019), the presence of the WRKY binding site led to repressed expression in the light (Fig. 5f). These trends were confirmed by mutational analysis. Mutations that disrupt a binding site for WRKY transcription factors increased the light-dependency of the promoter, while mutations that disrupt a binding sites for TCP transcription factors led to a noticeable, albeit not significant, decrease in light-dependency (Supplementary Fig. 15).

### Design of synthetic plant promoters

After identifying key features of native plant promoters, we sought to use these features in the design of synthetic promoters. We started by generating random sequences with nucleotide frequencies resembling either an average Arabidopsis or average maize promoter (Fig. 6a). We designed 10 sequences each for the two nucleotide frequencies; however, due to their AT-rich nature, the synthesis of approximately half of the sequences with an Arabidopsis promoter-like base composition failed. Consistent with the findings for native promoters, the synthetic promoters with low GC content, similar to that of Arabidopsis promoters, were 30% more active in tobacco leaves than those with GC content similar to that of maize promoters (Fig. 6b,c). However, as expected, these random synthetic promoters were weak. To increase their activity, we modified them by adding an Inr, Y patch element or TATA-box (Fig. 6a). Although all three of these core promoter elements, both alone and in combination, increased promoter strength, the TATA-box showed the strongest effect and the Inr the weakest (Fig. 6b,c and Supplementary Table 6). The relative activity of these three elements was similar across synthetic promoters with initial nucleotide frequencies similar to either Arabidopsis or maize and across the two assay systems. However, in tobacco leaves, the absolute change in promoter strength was different for synthetic promoters of different GC content, indicating that the elements tested in this assay system require a favorable sequence environment to achieve full activity (Fig. 6b). Taken together, the results demonstrate that it is possible to rationally design synthetic core promoters of varying strength by choosing an appropriate background nucleotide frequency and adding canonical core promoter elements. The strongest synthetic promoters reached activities comparable to the viral 35S minimal promoter.

**Fig. 6.**
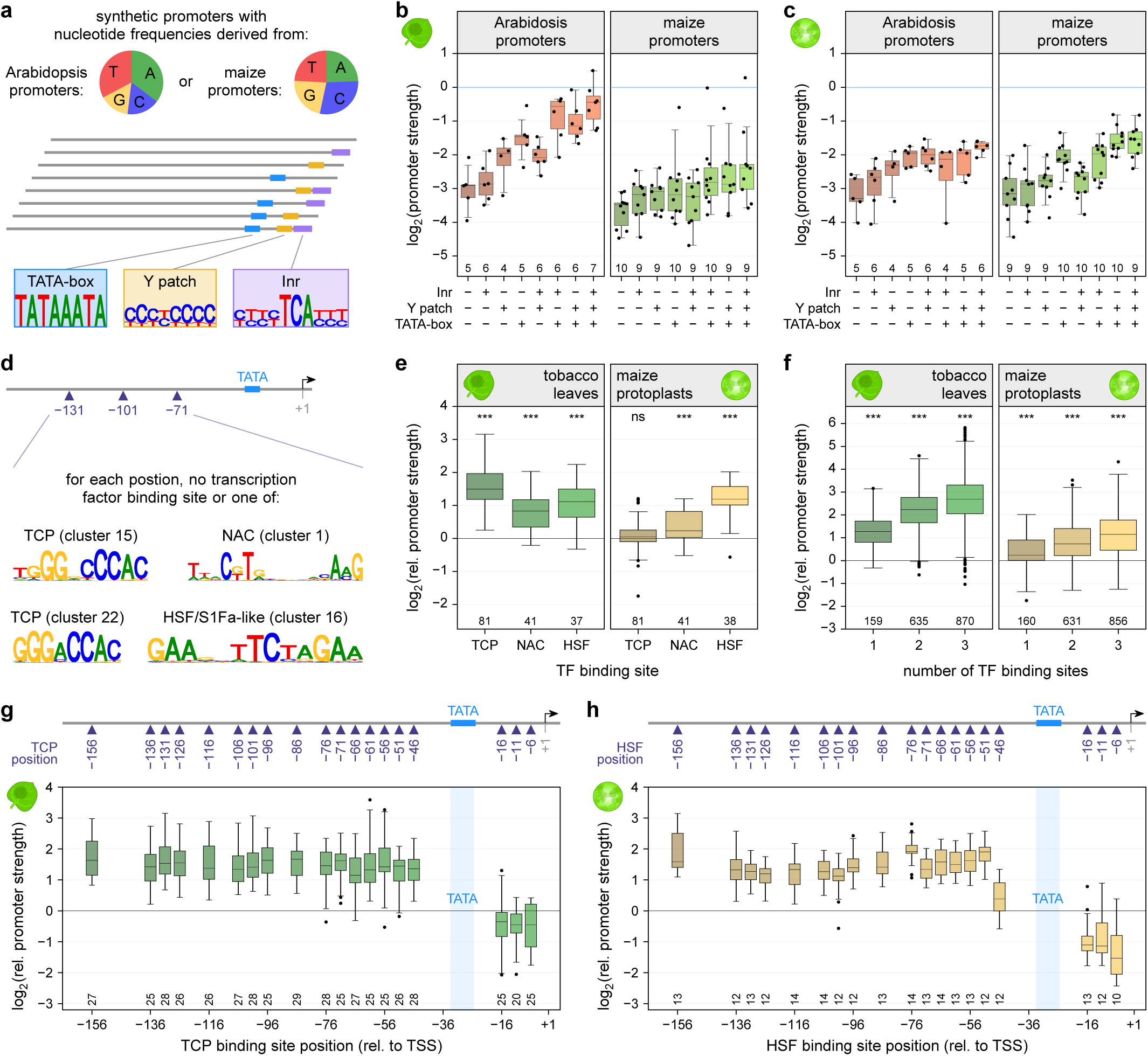
Design and validation of synthetic promoters. **a**-**c**, Synthetic promoters with nucleotide frequencies similar to an average Arabidopsis (35.2% A, 16.6% C, 15.3% G, 32.8% T) or maize (24.5% A, 29.0% C, 22.5% G, 23.9% T) promoter were created and modified by adding a TATA-box, Y patch, and/or Inr element (**a**). Promoter strength was determined by STARR-seq in tobacco leaves (**b**) and maize protoplasts (**c**). Promoters with an Arabidopsis-like nucleotide composition are shown on the left, those with maize-like base frequencies on the right. The strength of the 35S minimal promoter is indicated by a horizontal blue line. Due to the small sample size, individual data points are shown. **d**-**f**, Transcription factor binding sites for TCP, NAC, and HSF transcription factors were inserted at positions 35, 65, and/or 95 of the synthetic promoters with a TATA-box (**d**) and the activity of promoters with a single binding site for the indicated transcription factor (**e**) or multiple binding sites (**f**) was determined in tobacco leaves (left panel) or maize protoplasts (right panel). **g**,**h**, A single TCP (**g**) or HSF (**h**) transcription factor binding site was inserted at the indicated position in the synthetic promoters containing a TATA-box. The strength of these promoters was measured in tobacco leaves (**g**) or maize protoplasts (**h**). Boxplots are as defined in Figure 3. In (**e**-**h**), the corresponding promoter without any transcription factor binding was set to 0 (horizontal black line).

We also used the synthetic promoters to further analyze the effect of promoter-proximal TF binding sites. We focused on four different binding sites: two sites for TCP TFs, and one each for HSF TFs and NAC TFs. The TF binding sites were introduced at three positions in the synthetic promoters in which a TATA-box had been added (Fig. 6d). Because we did not observe position-dependent differences for any of the three TF binding sites, we grouped their respective data to perform the subsequent analyses. Consistent with our observations for native promoters, the TCP binding sites had the strongest effect in tobacco leaves, the HSF sites were most active in maize protoplasts, and the NAC sites had a weak but consistent effect across both assay systems (Fig. 6e). When more than one TF binding site was introduced into the synthetic promoters, their activities were additive, and the relative strengths of the promoters were conserved in combinations. The more binding sites that were present, the higher the promoter strength (Fig. 6f, Supplementary Fig. 16, and Supplementary Table 6).

Finally, to test whether the TFs show position-dependent activity with regard to the TATA-box, the binding sites for TCP, HSF, and NAC TFs were inserted at several positions upstream and downstream of the TATA-box. While these TF binding sites at all tested positions upstream of the TATA-box led to similar increases in promoter strength, they did not increase promoter strength when inserted downstream of the TATA-box (Fig. 6g,h, Supplementary Fig. 17, and Supplementary Table 6). These results likely reflect competition with the core transcriptional machinery that binds to this region.

**Fig. 7.**
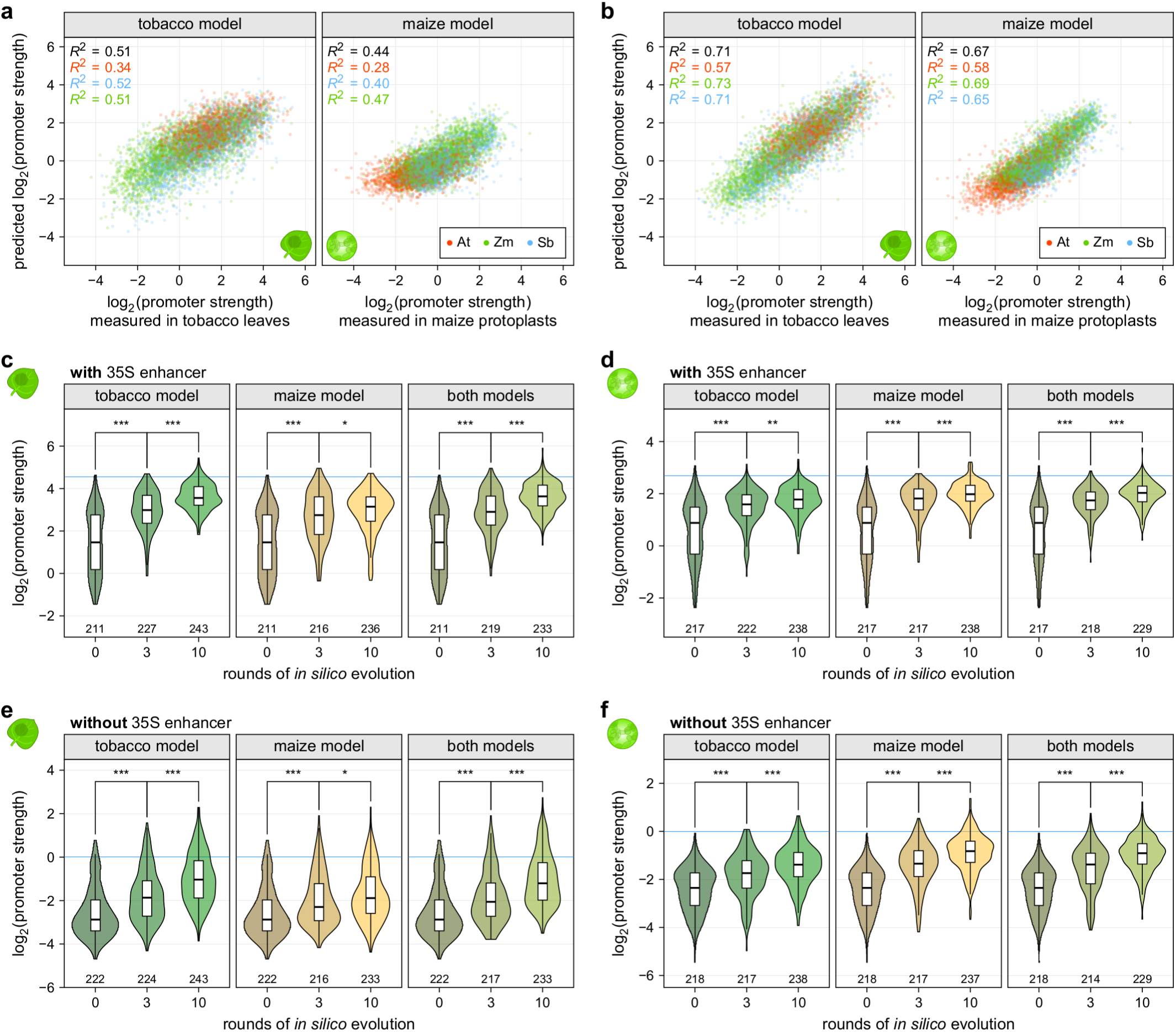
Computational models can predict promoter strength and enable *in silico* evolution of plant promoters. **a**, Correlation between the promoter strength as determined by STARR-seq using promoter libraries with the 35S enhancer and predictions from a linear model based on the GC content and motif scores for core promoter elements and transcription factors. The models were trained on data from the tobacco leaf system (tobacco model) or the maize protoplasts (maize model). The overall correlation is indicated in black and correlations for each species are colored as indicated (inset). Correlations (Pearson’s *R*^2^) are shown for a test set of 10% of all promoters. **b**, Similar to (**a**) but the prediction is based on a convolutional neural network trained on promoter sequences. **c**-**f**, Violin plots (as defined in Figure 1) of promoter strength of the unmodified promoters (0 rounds of evolution) or after they were subjected to three or ten rounds of *in silico* evolution as determined in tobacco leaves (**c,e**) or maize protoplasts (**d,f**). The promoters were tested in a library with (**c,d**) or without (**e,f**) an upstream 35S enhancer. The models used for the *in silico* evolution are indicated on each plot. The promoter strength of the 35S promoter is indicated by a horizontal blue line.

### Use of computational models to predict core promoter strength and improve promoter activity

Computational models have been employed to optimize synthetic gene-regulatory sequences (Cuperus et al., 2017; Kotopka and Smolke, 2020). Therefore, we set out to develop predictive models for core promoter strength using the data from the libraries with the 35S enhancer to train the models, as they had a better replicate correlation. For each assay system, we trained a separate model using 90% of the promoters, with the remaining 10% used to validate the model. We initially used a linear regression model for this task. The GC content and the maximum score for a match to the position weight matrices for the core promoter elements and TF clusters of each sequence were used as input features. The linear models explained 51% and 45% of the variability in promoter strength in tobacco leaves and maize protoplasts, respectively (Fig. 7a). In both systems, the TATA-box score was the most important feature for promoter strength, followed by GC content.

To obtain models with increased predictive power, we turned to a machine learning approach using a convolutional neural network (CNN). The models used the DNA sequence of the core promoters as input and predicted the strength of the promoters in the test set, resulting in an *R*^2^ of 0.71 and 0.67 for the tobacco and the maize systems, respectively (Fig. 7b).

We used these models for *in silico* evolution of 150 native promoters with weak, intermediate or strong activity in our assay. Additionally, we subjected the synthetic promoters with or without various core promoter elements to evolution. For each promoter, we generated every possible single nucleotide substitution variant and scored these variants with the CNN models. The best variant was retained and subjected to another round of evolution. We synthesized the starting sequences and those obtained after three and ten rounds of evolution and experimentally determined their activity. As predicted, we observed a large increase in promoter strength after three rounds of evolution and another, albeit less pronounced, increase after ten rounds (Fig. 7c,d and Supplementary Fig. 18). We obtained the best results when the evolution was performed with the CNN model trained on data from the same assay system. However, when we used a combination of both models to score the promoter variants, we could generate promoters with high activities in both systems that were on par with those evolved with the CNN model that was trained on data from the system in which the evolved sequences were tested (Fig. 7c-f and Supplementary Table 7). The models used for the *in silico* evolution were trained on data from libraries with an upstream 35S enhancer; however, when we tested the evolved promoters without the 35S enhancer, their activities followed the same trend, with a large increase in activity after three rounds and an additional increase after ten (Fig. 7e,f). These results suggest that the increased promoter strength generated by the evolution process was not enhancer-dependent and that these promoters might similarly work well with other enhancers.

## Discussion

The use of plants to synthesize medical and nutritional products requires precise control of foreign genes; similarly, precise control of endogenous genes is required to generate plants that can better withstand stresses. This precision can be realized through the design of synthetic promoters with optimal sequences, spacings and orientations of regulatory elements. Here, we used the STARR-seq assay to characterize plant core promoters in depth. We demonstrate that the most critical element of a strong plant core promoter is the presence of a TATA-box approximately 30 to 40 bp upstream of the TSS. The next most critical element is a nucleotide composition appropriate for the plant that is being engineered. A promoter can further be improved with an Inr motif at the TSS and a pyrimidine-rich region between the TATA-box and the Inr. Such rationally designed promoters can reach activities comparable to the highly active viral 35S minimal promoter.

While it might be optimal to conduct these experiments within the genomic context *in planta*, current technologies make such large-scale studies feasible only with transient expression of reporter constructs. However, the lack of genomic context may be less important for promoter strength than is commonly assumed. Studies in human and *Drosophila* cells found that results from plasmid-based regulatory elements are highly correlated with those from genome-integrated ones in massively parallel reporter assays (Arnold et al., 2013; Klein et al., 2020). Moreover, human core promoters retain their relative strength regardless of where they are inserted in the genome or if they drive expression of a plasmid-encoded reporter; the genomic context appears merely to scale their activity but does so independently of promoter identity (Hong and Cohen, 2021). Furthermore, we and others have previously demonstrated that transient STARR-seq assays in plants recapitulate the relative strength and the condition-specificity of known regulatory elements (Ricci et al., 2019; Jores et al., 2020). Our findings about the relative strength of promoters should, therefore, apply to promoters integrated in the genome, with the caveat that nearby enhancers may modulate the absolute expression level in addition to tissue- and condition-specificity.

Promoter activity and conditional response can be further modified by the addition of TF binding sites upstream of the TATA-box. Such binding sites affected promoter strength in an additive manner. The choice of binding site, however, will depend on the assay system and on the TFs that are present and active in it. TF presence and activity cannot simply be inferred from TF motifs because plant TF families are large and often encode both activating and repressing factors with highly similar binding preferences. However, single cell genomics can determine which TFs are expressed in specific cell types and associated with chromatin accessibility of regulatory elements (Dorrity et al., 2020; Marand et al., 2020). This knowledge offers a promising avenue to explore the activity of cell type-specific regulatory elements. In the absence of an assay system derived from a cognate cell type, cell type-specific transcription factors can be co-expressed in the assay systems used here. Alternatively, a large array of promoters can be designed with an assortment of TF binding sites, followed by an assay like the one described here to identify the most active ones.

Nevertheless, the design of strong core promoters appears feasible without such cell type-specific or even species-specific data. Our CNN models accurately predicted promoter strength and could be used for *in silico* evolution to yield native and synthetic promoters with increased activity. Moreover, a combination of CNN models trained on data from the tobacco and maize assay systems yielded promoters active in both systems. Such promoters are robust candidates to use across a broad range of tissues and species and in conjunction with multiple enhancers.

In animals, enhancer-promoter interactions are fine-tuned to execute distinct regulatory programs, like expression of housekeeping or developmental genes (Gehrig et al., 2009; Arnold et al., 2017). Here, we studied the effect of only the viral 35S enhancer on plant promoters. However, this assay could be applied to study interactions between promoters and native plant enhancers; such experiments might reveal specific interactions between distinct types of promoters and enhancers. Combining the potent core promoters characterized here with equally well-characterized enhancers will add the desired condition-specific and cell type-specific regulation needed for applications in plant engineering and biotechnology.

## Methods

### Library design and construction

For this study, we used the sequence from -165 to +5 relative to the annotated transcription start site as core promoters. We used the Araport11 annotation (Cheng et al., 2017) for Arabidopsis (*Arabidopsis thaliana* Col-0) and the NCBI_v3.43 annotation (McCormick et al., 2018) for sorghum (*Sorghum Bicolor* BTx623). For maize (*Zea mays* L. cultivar B73) promoters, we used experimentally determined TSSs (Mejía-Guerra et al., 2015) and supplemented this set with the B73_RefGen_v4.42 annotation (Jiao et al., 2017) for genes without an experimentally confirmed TSS. The core promoter sequences were ordered as an oligo pool from Twist Biosciences.

The STARR-seq plasmids used herein are based on the plasmid pPSup (https://www.addgene.org/149416/; Jores et al., 2020). It harbors a phosphinothricin resistance gene (BlpR) and a GFP reporter construct terminated by the polyA site of the *Arabidopsis thaliana* ribulose bisphosphate carboxylase small chain 1A gene in the T-DNA region. The plant core promoters followed by a 5′ UTR from maize (Zm00001d041672; used for the Arabidopsis, maize and validation promoter libraries) or sorghum (SORBI_3010G047100; used the sorghum promoter library) histone H3 gene, an ATG start codon and a 12 bp random barcode (VNNVNNVNNVNN; V = A, C, or G) was cloned in front of the second codon of GFP by Golden Gate cloning (Engler et al., 2008). For control constructs, the 35S minimal promoter was used instead of the plant core promoters. Each library was bottlenecked to contain, on average, 10 to 20 barcodes per promoter. The 35S core was inserted upstream of the core promoters by Golden Gate cloning. The sequences of the 5′ UTRs and the 35S enhancer and minimal promoter are listed in Supplementary Table 8. All primers are listed in Supplementary Table 9. The STARR-seq plasmid libraries were introduced into *Agrobacterium tumefaciens* GV3101 strain harboring the helper plasmid pSoup (Hellens et al., 2000) by electroporation.

### Tobacco cultivation and transformation

Tobacco (*Nicotiana benthamiana*) was grown in soil (Sunshine Mix #4) at 25 °C in a long-day photoperiod (16 h light and 8 h dark; cool-white fluorescent lights [Philips TL-D 58W/840]; intensity 300 μmol/m^2^/s). Plants were transformed 3 to 4 weeks after germination. For transformation, an overnight culture of *Agrobacterium tumefaciens* was diluted into 100 mL YEP medium (1% [w/v] yeast extract, 2% [w/v] peptone) and grown at 28 °C to an OD of approximately 1. A 5 mL input sample of the cells was taken, and plasmids were isolated from it. The remaining cells were harvested and resuspended in 100 mL induction medium (M9 medium supplemented with 1% [w/v] glucose, 10 mM MES, pH 5.2, 100 μM CaCl_2_, 2 mM MgSO_4_, and 100 μM acetosyringone). After overnight growth, the bacteria were harvested, resuspended in infiltration solution (10 mM MES, pH 5.2, 10 mM MgCl_2_, 150 μM acetosyringone, and 5 μM lipoic acid) to an OD of 1 and infiltrated into the first two mature leaves of 3 to 6 tobacco plants. The plants were further grown for 48 h under normal conditions or in the dark prior to mRNA extraction.

### Maize protoplast generation and transformation

We used a slightly modified version of a published protoplasting and electroporation protocol (Sheen, 1990). Maize (*Zea mays* L. cultivar B73) seeds were germinated for 4 days in the light and the seedlings were grown in soil at 25 °C in the dark for 9 days. The center 8 to 10 cm of the second leaf from 10 to 12 plants were cut into thin strips perpendicular to the veins and immediately submerged in 10 mL protoplasting solution (0.6 M mannitol, 10 mM MES, 15 mg/mL cellulase R-10 [GoldBio], 3 mg/mL Macerozyme R-10 [GoldBio], 1 mM CaCl_2_, 5 mM β-mercaptoethanol, 0.1% [w/v] BSA, pH 5.7). The mixture was covered to keep out light, vacuum infiltrated for 30 min, and incubated with 40 rpm shaking for 2 h. Protoplasts were released with 80 rpm shaking for 5 min and filtered through a 40 μm filter. The protoplasts were harvested by centrifugation (3 min at 200 x g, room temperature) in a round bottom glass tube and washed with 3 mL ice cold electroporation solution (0.6 M mannitol, 4 mM MES, 20 mM KCl, pH 5.7). After centrifugation (2 min at 200 x g, room temperature), the cells were resuspended in 3 mL ice cold electroporation solution and counted. Approximately 1 million cells were mixed with 25 μ L, transferred to a 4 mm electroporation cuvette and incubated for 5 min on ice. The cells were electroporated (300 V, 25 µFD, 400 Ω) and 900 μL ice cold incubation buffer (0.6 M mannitol, 4 mM MES, 4 mM KCL, pH 5.7) was added. After 10 min incubation on ice, the cells were further diluted with 1.2 mL incubation buffer and kept at 25 °C in the dark for 16 h before mRNA collection. To cover each library, four electroporation reactions were performed, except for the smaller validation libraries in which two electroporation reactions were performed. For the maize protoplast STARR-seq, the plasmid library used for electroporation was sequenced as the input sample.

### STARR-seq assay

For each STARR-seq experiment, two independent biological replicates were performed. Different plants and fresh *Agrobacterium* cultures were used for each biological replicate and the replicates were performed on different days. For experiments in tobacco, 12 transformed leaves were collected from 6 plants. They were frozen in liquid nitrogen, ground in a mortar and immediately resuspended in 25 mL TRIzol (Thermo Fisher Scientific). The suspension was cleared by centrifugation (5 min at 4,000 x g, 4 °C), and the supernatant was thoroughly mixed with 5 mL chloroform. After centrifugation (15 min at 4,000 x g, 4 °C), the upper, aqueous phase was transferred to a new tube, mixed with 5 mL chloroform and centrifuged again (15 min at 4,000 x g, 4 °C). 13 mL of the upper, aqueous phase was transferred to new tubes, and RNA was precipitated with 1.3 mL 8 M LiCl and 32.5 mL 100% (v/v) ethanol by incubation at –80 °C for 15 min. The RNA was pelleted (30 min at 4,000 x g, 4 °C), washed with 10 mL 70% (v/v) ethanol, centrifuged again (5 min at 4,000 x g, 4 °C), and resuspended in 1.5 mL nuclease-free water. The solution was split into two halves and mRNAs were isolated from each using 150 μL magnetic Oligo(dT)_25_ beads (NEB) according to the manufacturer’s protocol. The mRNAs were eluted in 40 μL. The two samples per library were pooled and supplemented with 10 μL DNase I buffer, 10 μL 100 mM MnCl_2_, 2 µL DNase I (Thermo Fisher Scientific), and 1 µL RNaseOUT (Thermo Fisher Scientific). After 1 h incubation at 37 °C, 2 μL 20 mg/mL glycogen (Thermo Fisher Scientific), 10 μL 8 M LiCl, and 250 μL 100% (v/v) ethanol were added to the samples. Following precipitation at –80 °C, centrifugation (30 min at 20,000 x g, 4 °C), and washing with 200 μL 70% (v/v) ethanol (5 min at 20,000 x g, 4 °C), the pellet was resuspended in 100 μL nuclease-free water. Eight reactions with 5 μL mRNA each and a GFP construct-specific primer were prepared for cDNA synthesis using SuperScript IV reverse transcriptase (Thermo Fisher Scientific) according to the manufacturer’s instructions. Half of the reactions were used as no reverse transcription control, in which the enzyme was replaced with water. After cDNA synthesis, the reactions were pooled and purified with DNA Clean & Concentrator-5 columns (Zymo Research). The barcode region was amplified with 10 to 20 cycles of PCR and read out by next generation sequencing. For the smaller validation libraries, only 6 leaves were used and all volumes except the reverse transcription were halved.

For the STARR-seq assay in maize protoplasts, transformed protoplasts were harvested by centrifugation (3 min at 200 x g, 4 °C) 16 h after electroporation. The protoplasts were washed three times with 1 mL incubation buffer and centrifuged for 2 min at 200 x g and 4 °C. The cells were resuspended in 600 μL TRIzol (Thermo Fisher Scientific) and incubated for 5 min at room temperature. The suspension was thoroughly mixed with 120 μL chloroform and centrifuged (15 min at 20,000 x g, 4 °C). The upper, aqueous phase was transferred to a new tube, mixed with 120 μL chloroform and centrifuged again (15 min at 20,000 x g, 4 °C). RNA was precipitated from 400 μL of the supernatant with 1 μL 20 mg/mL glycogen (Thermo Fisher Scientific), 40 μL 8 M LiCl, and 1 mL 100% (v/v) ethanol by incubation at –80 °C for 15 min. After centrifugation (30 min at 20,000 x g, 4 °C), the pellet was washed with 200 μL 70% (v/v) ethanol, centrifuged again (5 min at 20,000 x g, 4 °C), and resuspended in 200 μL nuclease-free water. mRNAs were isolated from this solution using 50 μ magnetic Oligo(dT)_25_ beads (NEB) according to the manufacturer’s protocol, and the mRNAs were eluted in 40 μ water. DNase I treatment and precipitation were performed as for the mRNAs obtained from tobacco plants but with half the volume. Reverse transcription, purification, PCR amplification and sequencing were performed as for the tobacco samples.

### Subassembly and barcode sequencing

Paired-end sequencing on an Illumnia NextSeq 550 system was used for the subassembly of promoters with their corresponding barcodes. The promoter region was sequenced using partially overlapping, paired 144 bp reads, and two 15 bp indexing reads were used to sequence the barcodes. The promoter and barcode reads were assembled using PANDAseq (Masella et al., 2012) and the promoters were aligned to the designed core promoter sequences. Promoter-barcode pairs with less than 5 reads and promoters with a mutation or truncation were discarded. Barcode sequencing was performed using paired-end reads on a Illumnia NextSeq 550 platform. The reads were trimmed to only the barcode portion assembled with PANDAseq. All sequencing results were deposited in the NCBI Sequence Read Archive under the BioProject accession PRJNA714258 (http://www.ncbi.nlm.nih.gov/bioproject/714258). The scripts used for processing the raw reads are available at https://github.com/tobjores/Synthetic-Promoter-Designs-Enabled-by-a-Comprehensive-Analysis-of-Plant-Core-Promoters.

### Computational methods

For analysis of the STARR-seq experiments, the reads for each barcode were counted in the input and cDNA samples. Barcode counts below 5 were discarded. Barcode enrichment was calculated by dividing the barcode frequency (barcode counts divided by all counts) in the cDNA sample by that in the input sample. The enrichment of the promoters was calculated as the median enrichment of all barcodes linked to them. We calculated the promoter strength as the log_2_ of the promoter enrichment normalized to the enrichment of 35S minimal promoter. We used the average promoter strength from both replicates for all analyses. Spearman and Pearson correlations were calculated using the base R function. Significance was determined using the two-sided Wilcoxon rank-sum test as implemented in base R. GO-term enrichment analysis was performed using the ggprofiler2 (version 0.1.9) (Raudvere et al., 2019) library for R and a custom gmt file with GOslim terms. Gene expression data was obtained from the EMBL-EBI Expression Atlas (https://www.ebi.ac.uk/gxa/about.html) using experiments E-MTAB-7978 (Mergner et al., 2020), E-GEOD-50191 (Walley et al., 2016), and E-MTAB-5956 (Wang et al., 2018) for Arabidopsis, maize, and sorghum, respectively. The tissue specificity index τ was calculated as previously published (Yanai et al., 2005). Sequences for TFIIB proteins were obtained from Uniprot (https://www.uniprot.org/; see Supplementary Table 10 for accession numbers) and aligned using Clustal Omega (Madeira et al., 2019). The code used for analyses is available at https://github.com/tobjores/Synthetic-Promoter-Designs-Enabled-by-a-Comprehensive-Analysis-of-Plant-Core-Promoters.

### Prediction of core promoter elements and transcription factor binding sites

The TATA-box and Inr motifs were obtained from the plant promoter database (Shahmuradov et al., 2003) and for each a consensus motif was created by merging the motifs from dicot and monocot promoters using the universalmotif (version 1.6.3) library for R. Motifs for BREu and BREd were obtained from JASPAR (Fornes et al., 2020). The motifs for the polypyrimidine initiator TCT and the Y patch were created from published sequences of these elements (Yamamoto et al., 2007; Parry et al., 2010). Binding site motifs for Arabidopsis transcription factors were obtained from the PlantTFDB (Tian et al., 2020). Transcription factor motifs were clustered by similarity using the compare_motifs() function from the R library universalmotif. The original clusters were improved by manual inspection and reannotation. Consensus motifs for the final transcription factor motifs were created using the merge_motifs() function from universalmotif. Meme files with the motifs used in this study are available at https://github.com/tobjores/Synthetic-Promoter-Designs-Enabled-by-a-Comprehensive-Analysis-of-Plant-Core-Promoters. Promoter sequences were analyzed with the universalmotif library assuming a neutral background nucleotide frequency. For the initiator elements, only the last 10 (Inr) or the last 6 (TCT) bases were scanned. For BREu and BREd, the sequences immediately upstream and downstream of highest scoring TATA-box were analyzed. For each sequence, the maximum motif score was calculated and normalized to the minimum (set to 0) and maximum (set to 1) scores possible. Sequences with a score of at least 0.85 were considered positive. For testing the effect of the BREu and BREd motifs (Supplementary Fig. 6), only sequences with a TATA-box score of at least 0.7 were considered.

### Design of validation sequences

To directly validate the importance of the TATA-box, BRE^u^, and BRE^d^ elements, we picked 30 promoters (10 each from Arabidopsis, maize, sorghum if possible) according to the following critera: for mutations of a canonical TATA-box, we selected promoters with a TATA-box motif score above 0.9 in the -59 to -23 region. The two conserved T nucleotides in the core TATA motif were replaced individually or together with Gs. We also selected 30 promoters with a maximum TATA-box motif score of 0.7 to 0.75. This weak TATA-box was replaced with either a canonical TATA-box motif (TATAAAT) or a mutated version of it (TAGAAAT). For the BRE elements, we first filtered for promoters with a TATA-box motif score of at least 0.85 in the -59 to -23 region. From these, we picked promoters with a BRE motif score above 0.85. For the BRE^u^ element, we mutated bases 3, 6, and 7 to T, A, and A respectively. For the BRE^d^ element, we mutated bases 2,4, and 6 to A. We also selected promoters where both the BRE^u^ and the BRE^d^ motif scores are below 0.5 to insert either a canonical BRE^u^ (AGCGCGCC) or BRE^d^ (GTTTGTT) element.

### Synthetic promoter design

Synthetic promoters were designed by generating 170 bp long random sequences with a nucleotide composition similar to an average Arabidopsis (35.2% A, 16.6% C, 15.3% G, 32.8% T) or maize (24.5% A, 29.0% C, 22.5% G, 23.9% T) promoter. We filtered out any random sequence with motif scores higher than 0.75 for a TATA-box, Inr, or Y patch element, or for transcription factor binding site of clusters 1, 15, 16, or 22. Promoters containing recognition sites for the restriction enzymes used for cloning (BsaI and BbsI) were also removed. From each set of promoters (Arabidopsis or maize nucleotide composition) that passed the filters, we randomly selected 10 variants for further modification. The promoters were kept as is, or modified with a TATA-box (TATAAATA) at positions 133 to 140, a Y patch (A and G nucleotides of the promoter were changed to C) at positions 147 to 154, and/or an Inr element (yyyyTCAyyy, where y indicates a change of A to T or G to C) at positions 160 to 169. To study the effect of transcription factors, the synthetic promoters with the TATA-box were chosen as backgrounds. Binding sites for NAC (cluster 1, TTACGTGnnnnACAAG, where n represents bases of the promoter background), TCP (cluster 15, TGGGGCCCAC and cluster 22, GGGACCAC), or HSF/S1Fa-like (cluster 16, GAAGCTTCTAGAA) transcription factors were inserted at various positions of these promoters.

### Computational modeling of promoter strength

To predict promoter strength, we built separate models for the tobacco leaf and the maize protoplast system. We used the results from the libraries with the 35S enhancer in the dark for training and validation. The models were trained on a set of 90% of all measured promoters and tested against the held-out set of the remaining 10% of the promoters.

We used the base R function lm() to build a linear model for predicting promoter strength based on the promoter’s GC content and its maximum motif score for six core promoter elements (TATA-box, Inr, TCT, BRE^u^, BRE^d^, and Y patch) and 72 consensus transcription factor binding motifs.

To build a direct sequence to promoter strength model we build a convolutional neural network using the tensorflow (version 2.2) package in python. The model consist of two forward- and reverse-sequence scan layers adapted from DeepGMAP (Onimaru et al., 2020) with 128 filters and a kernel width of 13 that feed into a regular convolutional layer (128 filters, kernel width 13, ReLU activation). Each convolutional layer is followed by a dropout layer with a 0.15 dropout rate. The output of the convolutional layers is fed into a dense layer with 64 filters with batch-normalization and ReLU activation that is followed by a final dense layer generating the single output. We initialized the first convolutional layer kernel with the clustered transcription factor motifs. The source code and the models are available on GITHUB.

### *In silico* evolution of promoter sequences

We used the convolutional neural networks to improve promoter performance in an iterative fashion. In each round, we generated all possible single nucleotide variants of a given promoter, scored them with the convolutional neural network models and kept the variant with the highest predicted activity for the next round. The sequences were scored with either just one of the models trained on the tobacco leaf or the maize protoplast data or with both models in which case the mean of both predictions was used to select the best performing variant. We experimentally tested these sequences after 3 and 10 rounds of this process. For the evolution, we selected native promoters showing either weak, intermediate, or strong activity in both assay systems or were strong in one system and weak in the other one. Additionally, we also performed the *in silico* evolution with the synthetic promoters described above.

### Data availability

All sequencing results are deposited in the NCBI Sequence Read Archive under the BioProject accession PRJNA714258 (http://www.ncbi.nlm.nih.gov/bioproject/714258).

## Code availability

The code used in this study is available on Github (https://github.com/tobjores/Synthetic-Promoter-Designs-Enabled-by-a-Comprehensive-Analysis-of-Plant-Core-Promoters).

## Acknowledgments

We thank Aimer Gutierrez Diaz and Erich Grotewold for providing maize TSS data. This work was supported by the National Science Foundation (RESEARCH-PGR grant 1748843 to E.S.B., S.F. and C.Q.), the German Research Foundation (DFG; fellowship 441540116 to T.J., and the National Institutes of Health (T32 training grant HG000035 to J.T.).

## Author contributions

All authors conceived and interpreted experiments and wrote the article; T.J. and J.T. performed experiments; and T.J. analyzed the data and prepared the figures. T.J. and T.W. did the *in silico* modelling.

## Competing Interests

The authors declare no competing interests.

---

**Supplementary Fig. 1.**
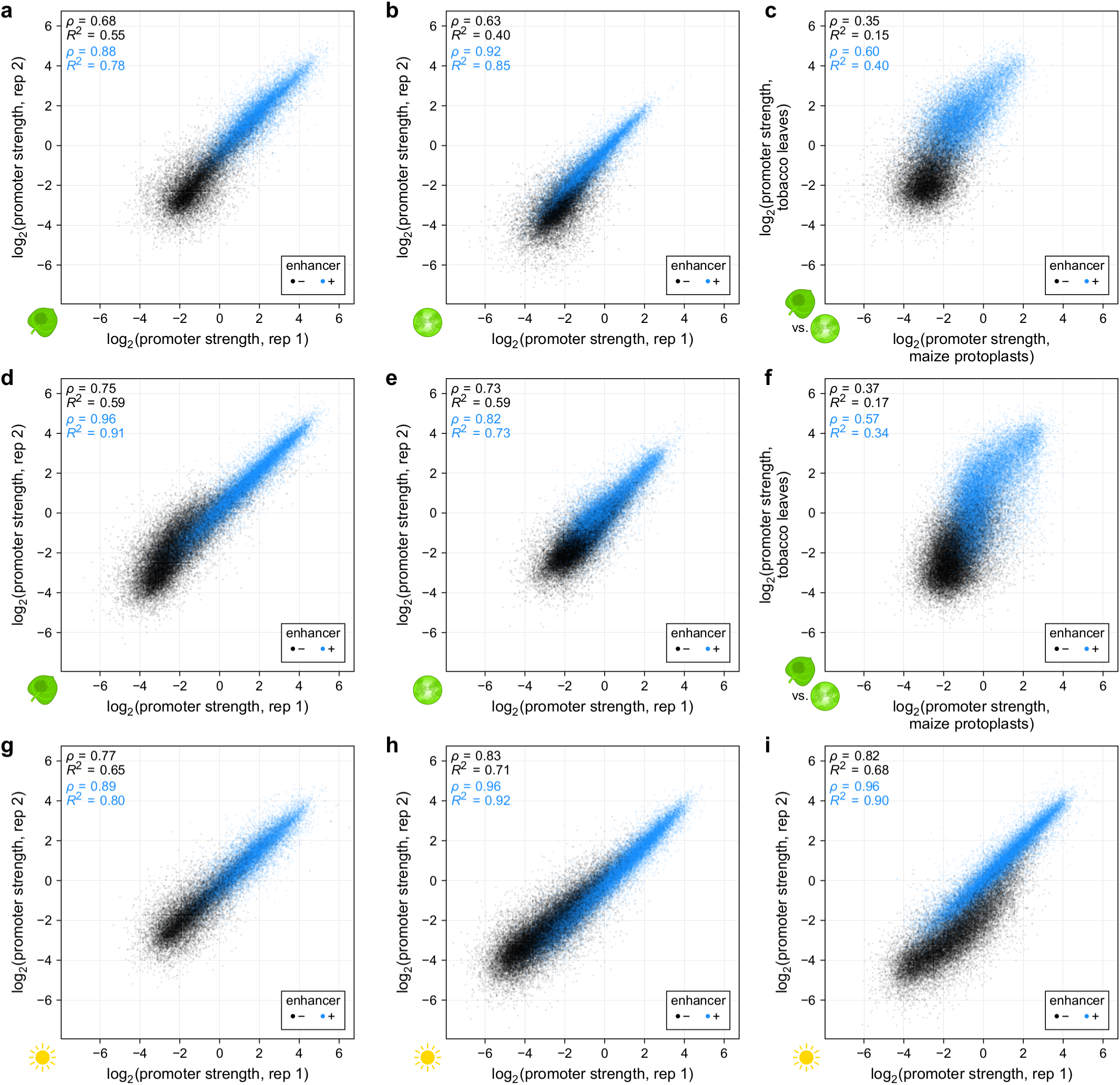
The promoter STARR-seq assay is highly reproducible but promoter strength depends on the assay system. **a**,**b**, Correlation of two biological replicates of STARR-seq using the Arabidopsis promoter libraries in tobacco leaves (**a**) or in maize protoplasts (**b**). **c**, Comparison of the strength of Arabidopsis promoters in tobacco leaves and maize protoplasts. **d**,**e**, Correlation of two biological replicates of STARR-seq using the sorghum promoter libraries in tobacco leaves (**d**) or in maize protoplasts (**e**). **f**, Comparison of the strength of sorghum promoters in tobacco leaves and maize protoplasts. **g**-**i**, Correlation of two biological replicates of STARR-seq using the Arabidopsis (**g**), maize (**h**), or sorghum (**i**) promoter libraries in tobacco leaves that were kept for two days in 16h light/8h dark cycles prior to mRNA extraction. Pearson’s *R*^2^ and Spearman’s *ρ* are indicated in all plots.

**Supplementary Fig. 2.**
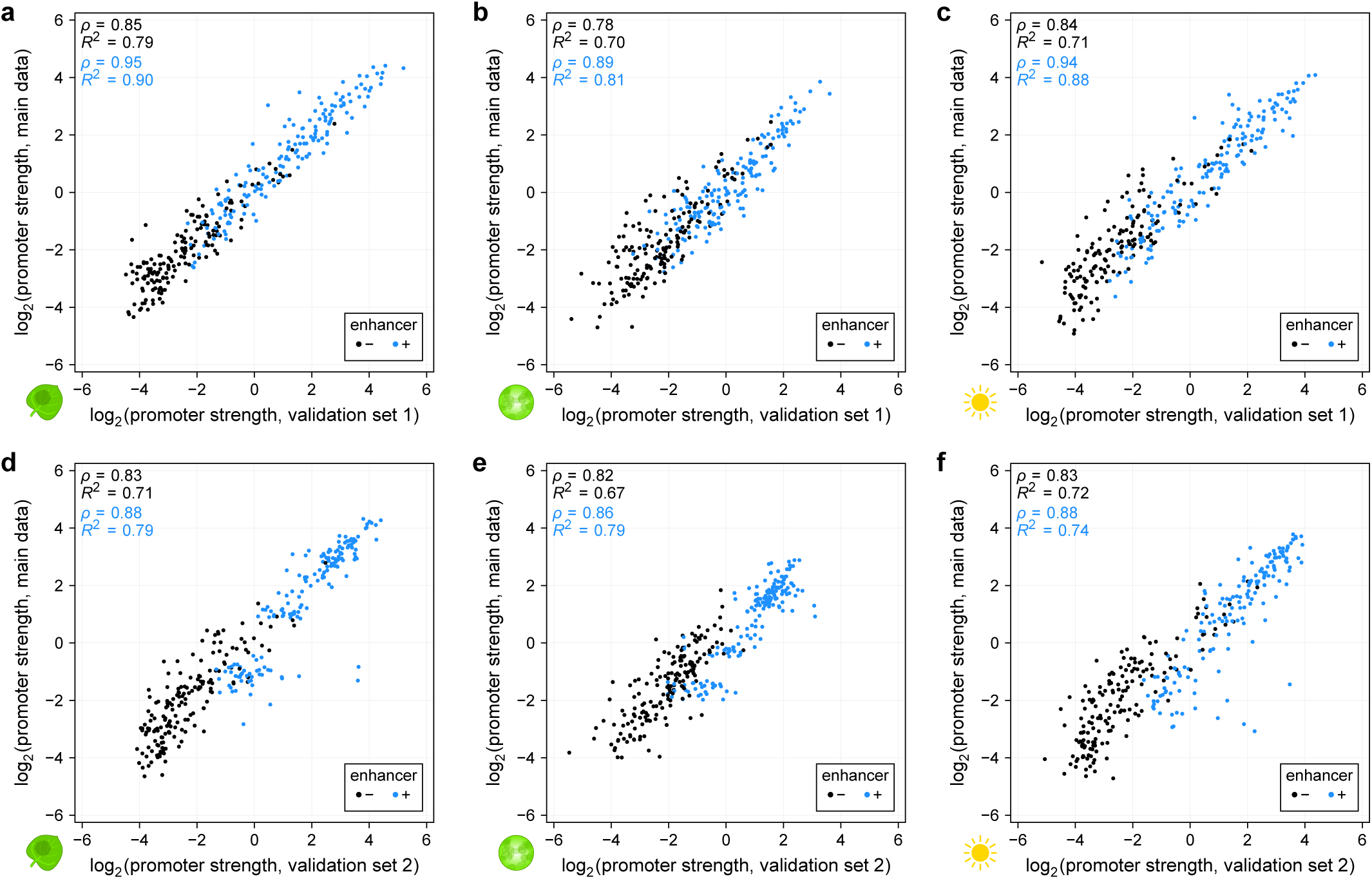
Promoter strength in small validation libraries correlates highly with comprehensive data. **a**-**c**, Correlation between the strength of promoters present in the comprehensive promoter libraries (main data) and in a separate, smaller validation library. The promoter strength was determined in tobacco leaves (**a**) and maize protoplasts (**b**) that were kept in the dark prior to mRNA extraction. Additionally, promoter strength was measured in tobacco leaves that were kept for two days in 16h light/8h dark cycles prior to mRNA extraction (**c**). **d**-**f**, As in (**a**-**c**) but for a second validation library. Pearson’s *R*^2^ and Spearman’s *ρ* are indicated in all plots.

**Supplementary Fig. 3.**
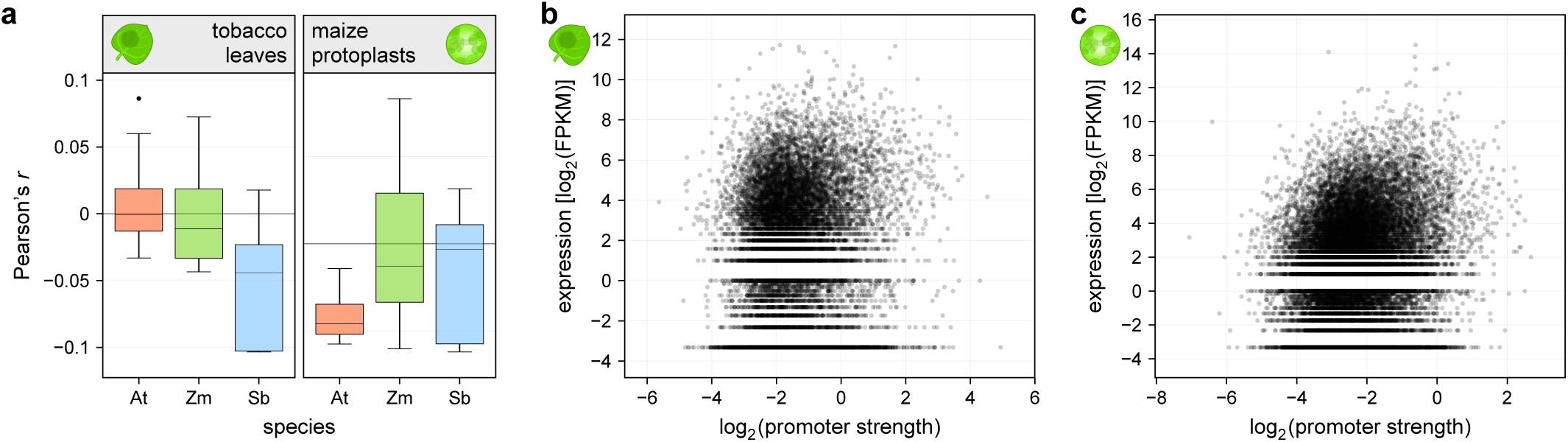
Promoter strength and *in vivo* expression levels are not correlated. **a**, Correlation (Pearson’s *r*) between the promoter strength and expression levels of the corresponding genes in the indicated species. Each boxplot (center line, median; box limits, upper and lower quartiles; whiskers, 1.5 × interquartile range; points, outliers) represents the correlation for all individual tissue samples in the RNA-seq dataset (see Methods). **b**,**c**, Examples of the correlation between gene expression (Arabidopsis adult cotyledon (**b**) or maize root cortex (**c**) samples) and promoter strength as determined in tobacco leaves (**b**) or maize protoplasts (**c**). These examples correspond to the highest correlations in (**a**).

**Supplementary Fig. 4.**
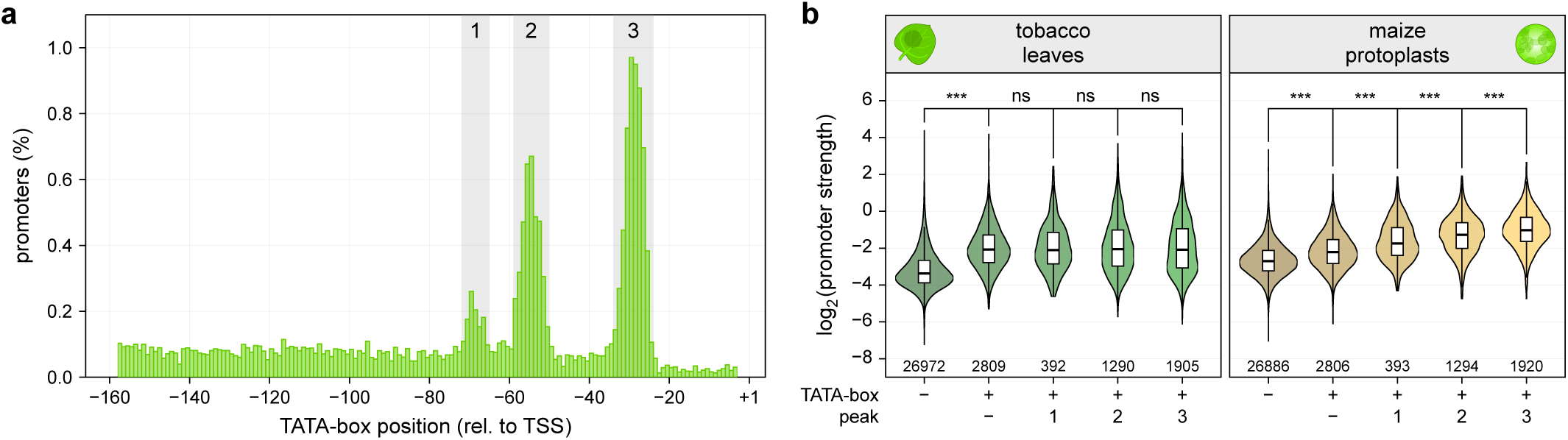
Strength of maize promoters depends on the TATA-box location in maize protoplasts. **a**, Histogram showing the percentage of maize promoters with a TATA-box at the indicated position (reproduced from Figure 3). Three peaks in the distribution of TATA-boxes are highlighted in gray. Peak 1 spans bases −72 to −65, peak 2 spans bases −59 to −50, and peak 3 spans bases −34 to −24. **b**, Violin plots (as defined in Figure 1) of promoter strength for maize promoters without enhancer in the indicated assay system. Promoters without a TATA-box (−) were compared to those with a TATA-box outside (+/−) or within one of the three peaks highlighted in (**a**).

**Supplementary Fig. 5.**
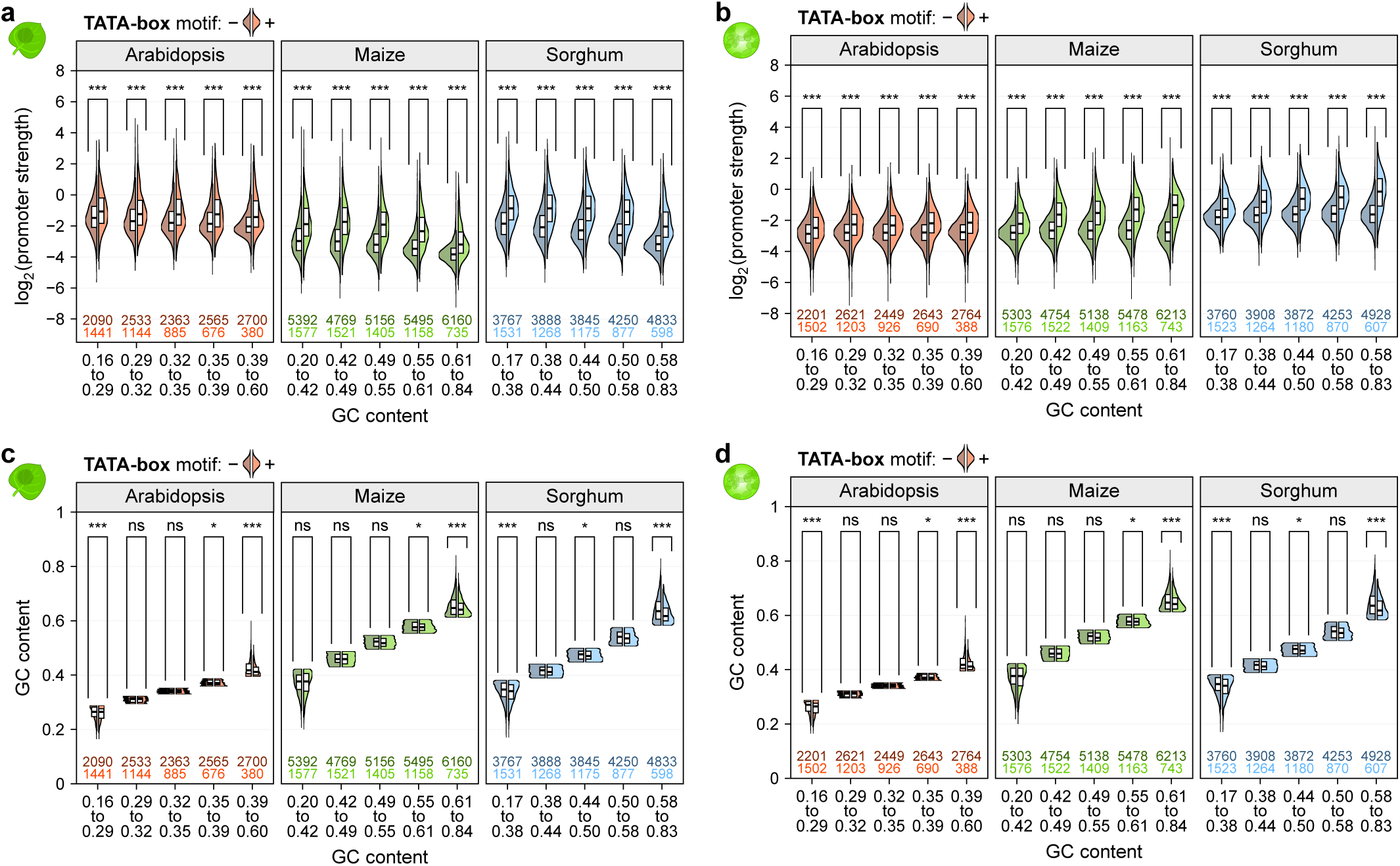
The effect of the TATA-box on promoter strength is not a result of decreased GC content. **a**-**d**, Violin plots of promoter strength (**a,b**) or GC content (**c,d**) in tobacco leaves (**a,c**) or maize protoplasts (**b,d**). Promoters were grouped by GC content and split into promoters without (left half, darker color) or with (right half, lighter color) a TATA-box. Violin plots are as defined in Figure 1, except only one half is shown.

**Supplementary Fig. 6.**
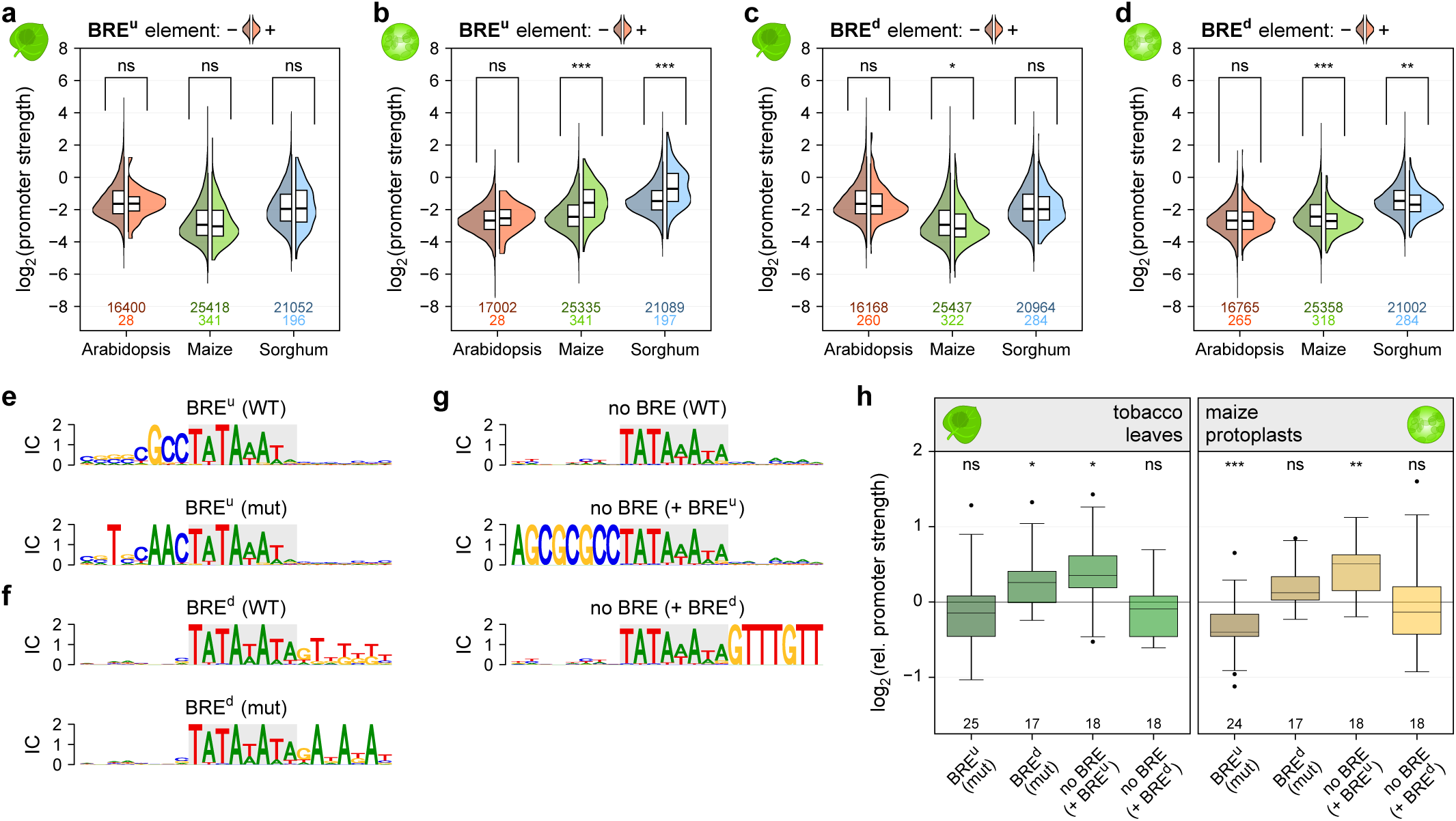
The BRE^u^ element is most active in maize protoplasts. **a**-**d**, Violin plots of promoter strength in tobacco leaves (**a,c**) or maize protoplasts (**b,d**). Promoters with a strong or intermediate TATA-box (motif score ≥ 0.7; see Methods) were grouped by GC content and split into promoters without (left half, darker color) or with (right half, lighter color) a BRE^u^ (**a,b**), or BRE^d^ (**c,d**) element. Violin plots are as defined in Figure 1, except only one half is shown. **e**,**g**, Logoplots for promoters with a BRE^u^ (**e**) or BRE^d^ (**f**) before (WT) and after (mut) introducing mutations that disrupt the elements. **g**, Logoplots for promoters without a BRE (WT) and with an inserted BRE^u^ (+ BRE^u^) or BRE^d^ (+ BRE^d^) element. **h**, Boxplots (as defined in Figure 3) for the relative strength of the promoter variants shown in (**e**-**g**). The corresponding WT promoter was set to 0 (horizontal black line).

**Supplementary Fig. 7.**
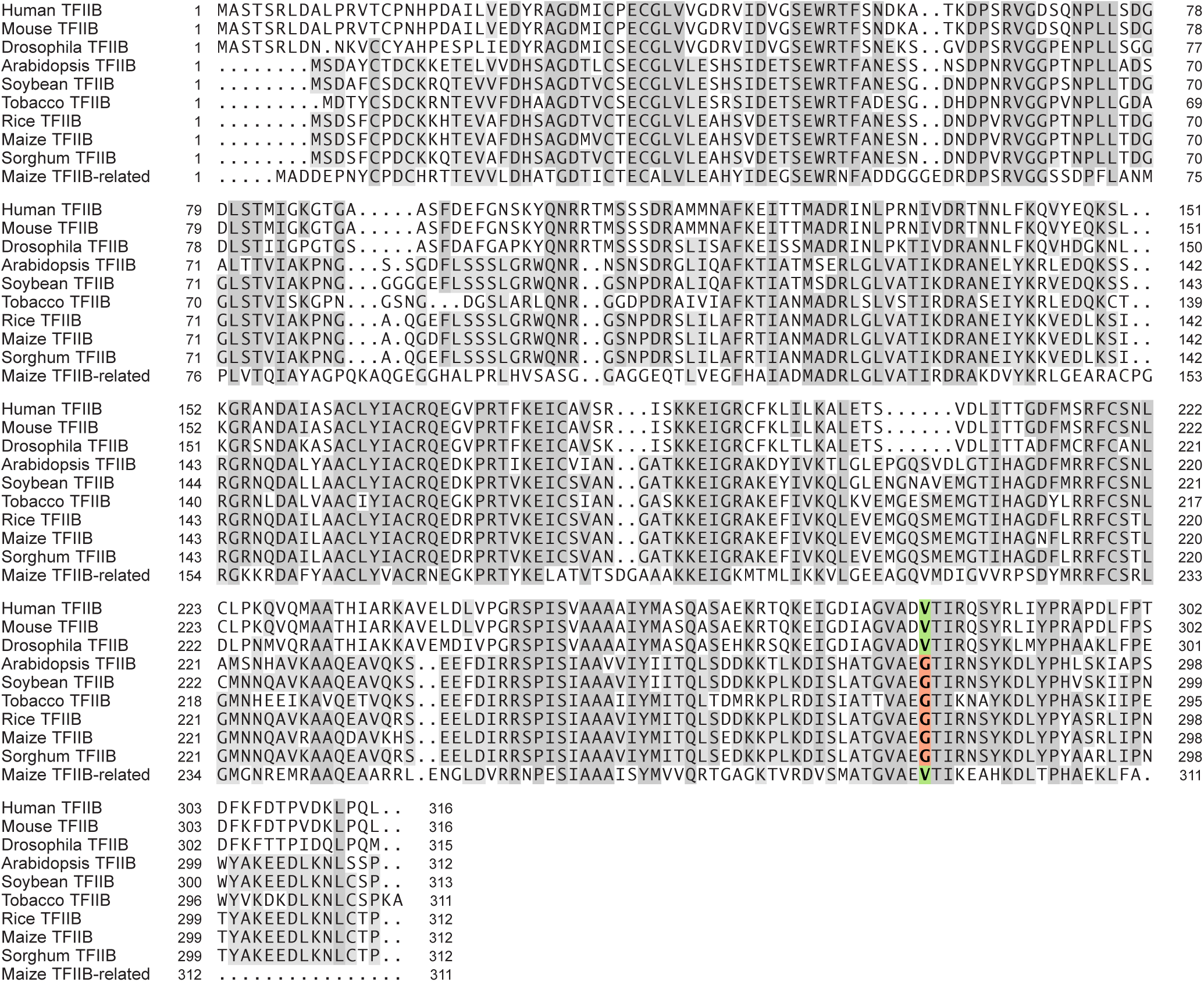
The maize genome encodes a TFIIB-related protein with a conserved valine residue required for BRE^u^ recognition. Alignment of TFIIB and TFIIB-like protein sequences from indicated species. Residues conserved in 80 or 50% of the sequences are highlighted in dark or light gray, respectively. The valine residue required for recognition of BRE^u^ is highlighted in green.

**Supplementary Fig. 8.**
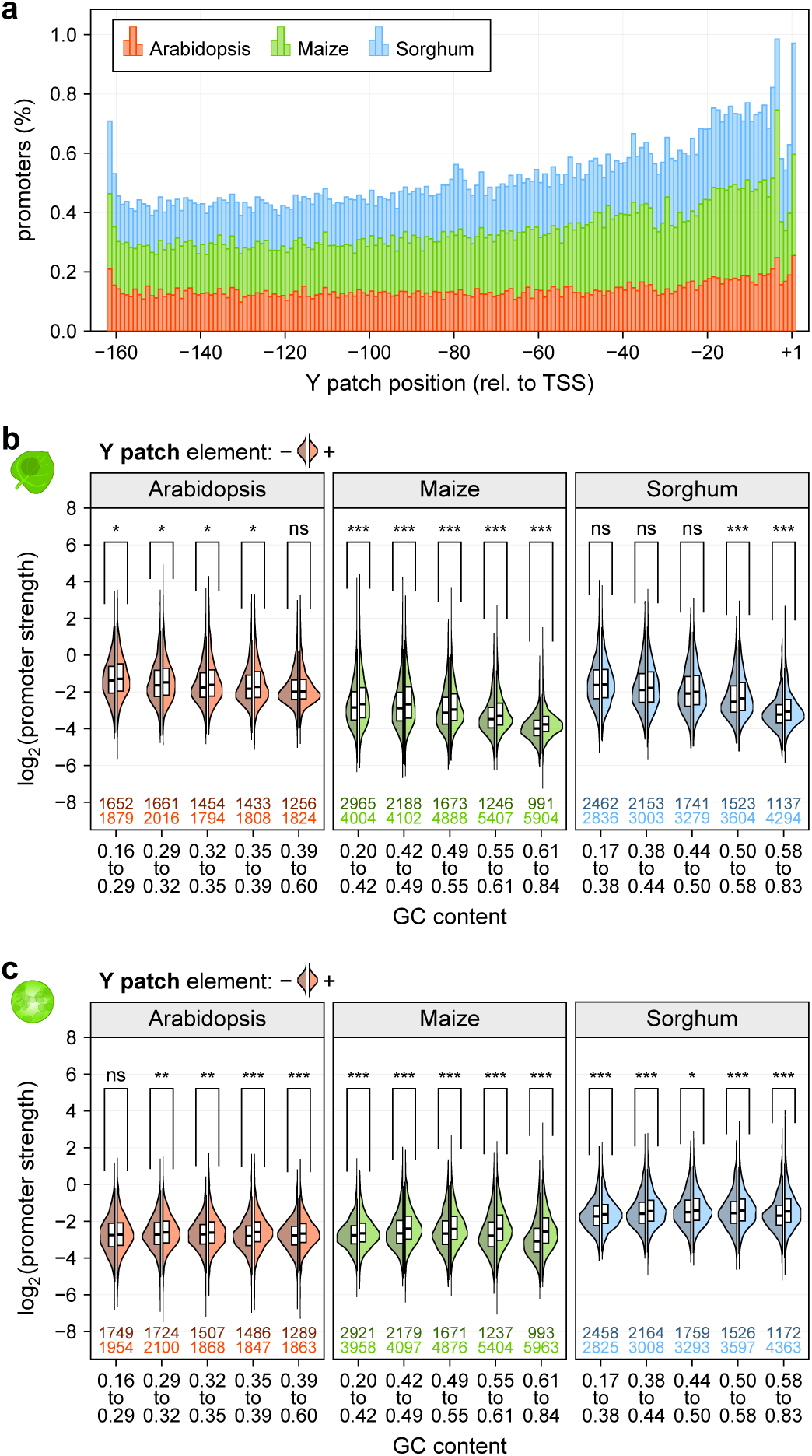
The Y patch is a plant-specific core promoter element. **a**, Histogram showing the percentage of promoters with a TATA-box at the indicated position. **b**,**c**, Violin plots of promoter strength in tobacco leaves (**b**) or maize protoplasts (**c**). Promoters were grouped by GC content and split into promoters without (left half, darker color) or with (right half, lighter color) a Y patch. Violin plots are as defined in Figure 1, except only one half is shown.

**Supplementary Fig. 9.**
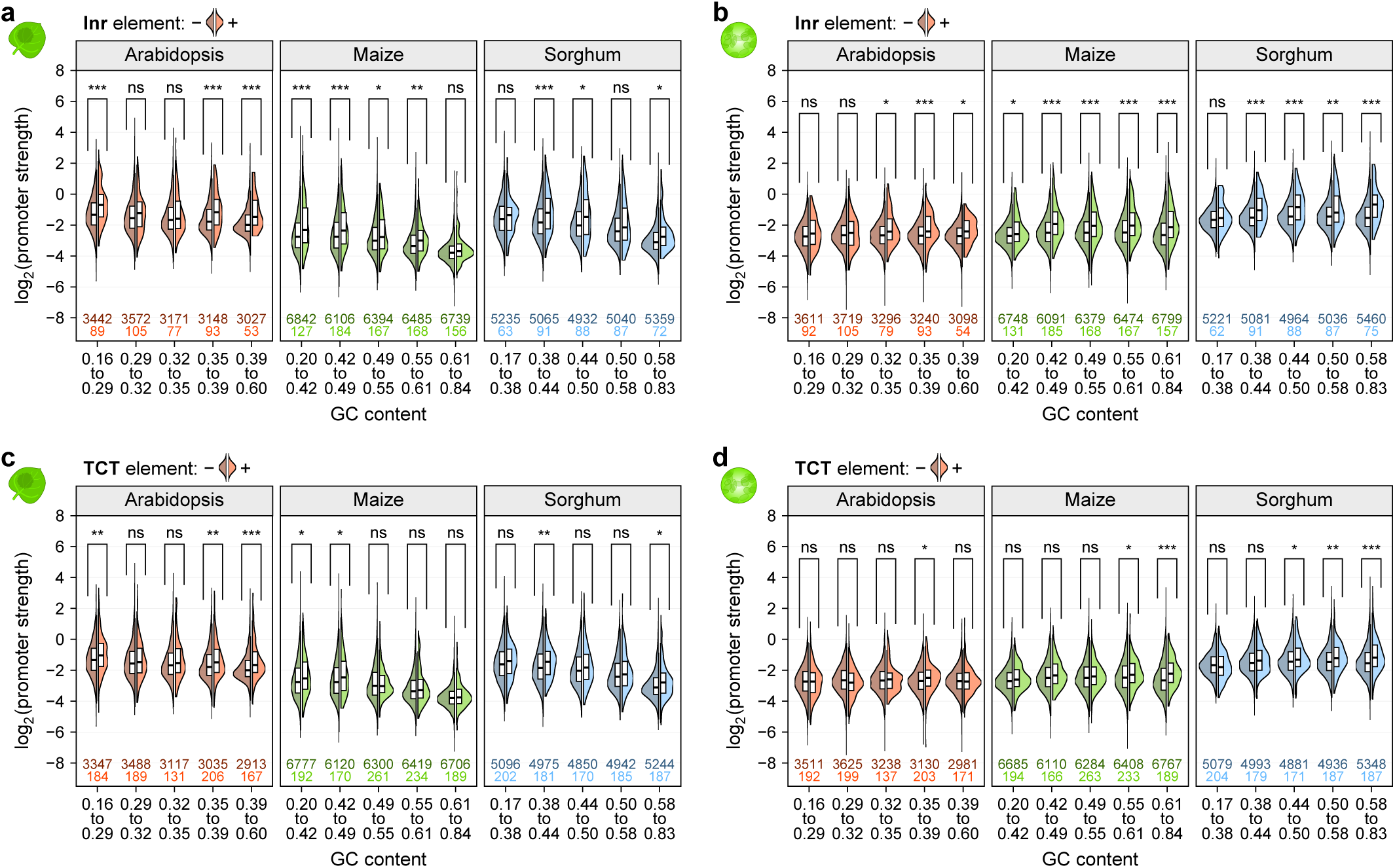
Core promoter elements at the TSS influence promoter strength. **a**-**d**, Violin plots of promoter strength in tobacco leaves (**a,c**) or maize protoplasts (**b,d**). Promoters were grouped by GC content and split into promoters without (left half, darker color) or with (right half, lighter color) an Inr (**a,b**), or TCT (**c,d**) element at the TSS. Violin plots are as defined in Figure 1, except only one half is shown.

**Supplementary Fig. 10.**
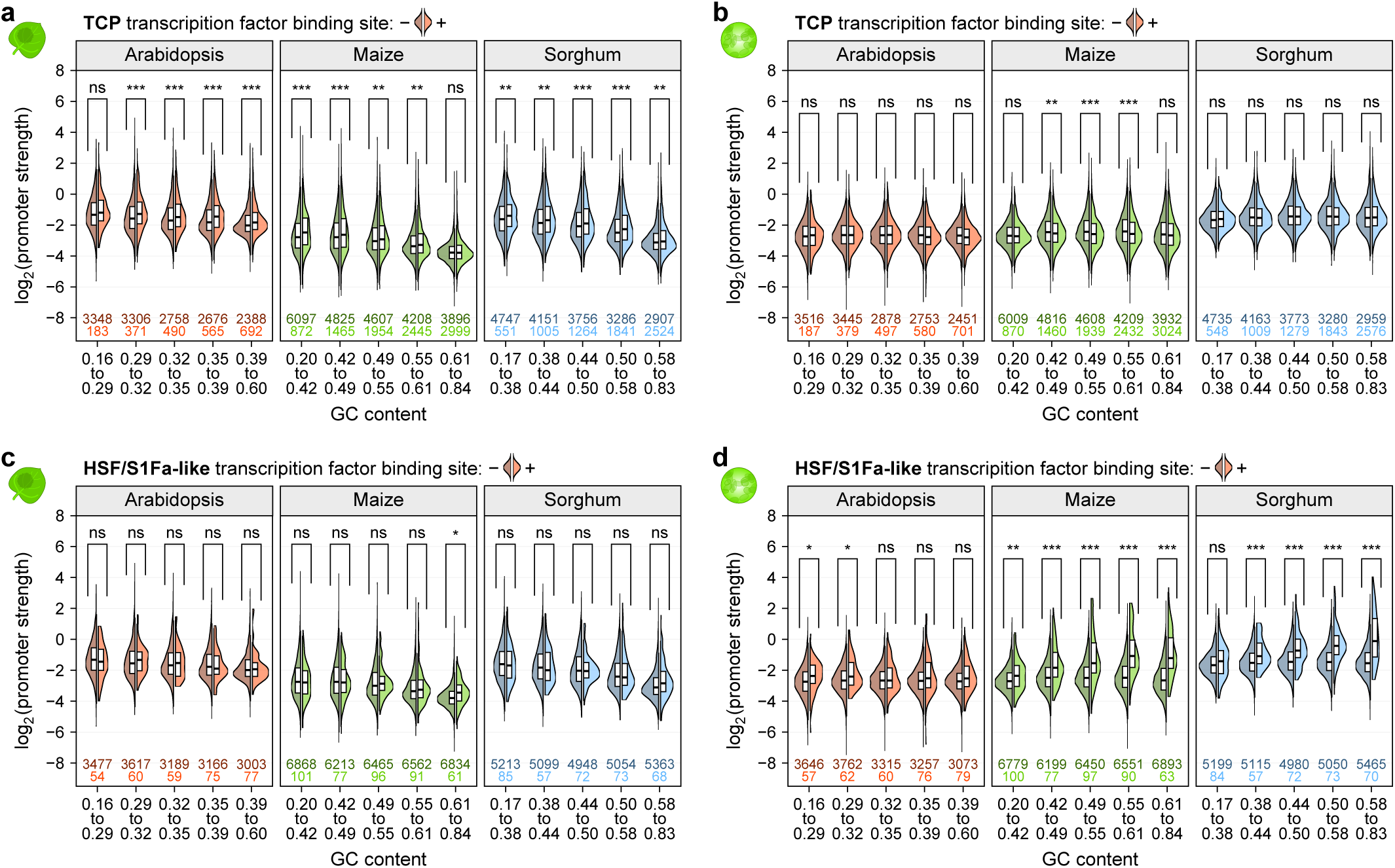
Transcription factor binding sites contribute to promoter strength in an assay system-dependent manner. **a**-**d**, Violin plots of promoter strength for libraries without enhancer in tobacco leaves (**a,c**) or maize protoplasts (**b,d**). Promoters were grouped by GC content and split into promoters without (left half, darker color) or with (right half, lighter color) a binding site for TCP (**a,b**) or HSF (**c,d**) transcription factors. Violin plots are as defined in Figure 1, except only one half is shown.

**Supplementary Fig. 11.**
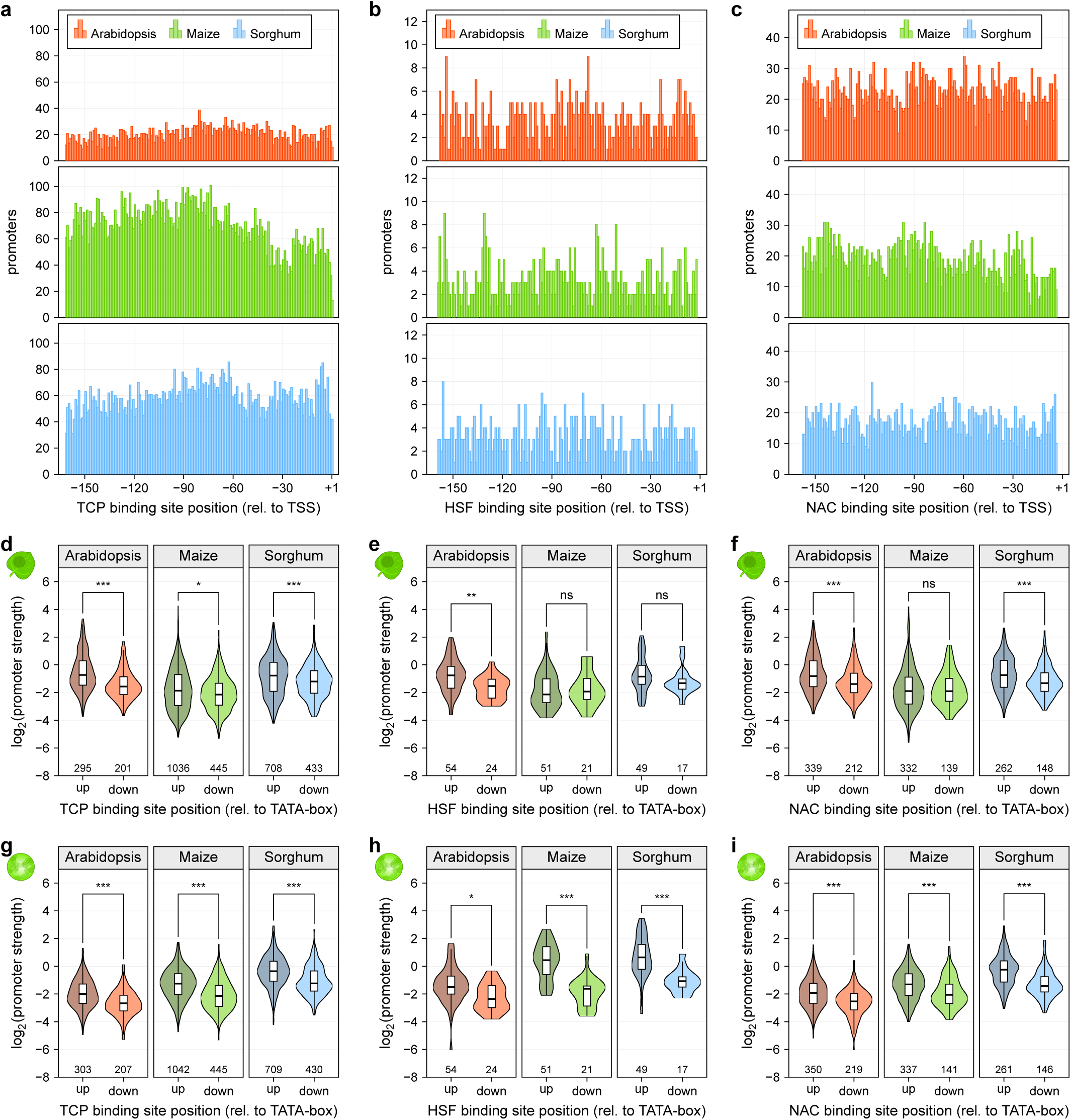
Transcription factor binding sites are more active upstream of the TATA-box. **a**-**c**, Histograms showing the number of promoters with a TCP (**a**), HSF (**b**) or NAC (**c**) transcription factor binding site at the indicated position. **d**-**i**, Violin plots (as defined in Figure 1) of promoter strength for libraries without enhancer in tobacco leaves (**d**-**f**) or maize protoplasts (**g**-**i**). Promoters were grouped by whether their TCP (**d,g**), HSF (**e,h**) or NAC (**f,i**) transcription factor binding site is located upstream (up) or downstream (down) of the TATA-box.

**Supplementary Fig. 12.**
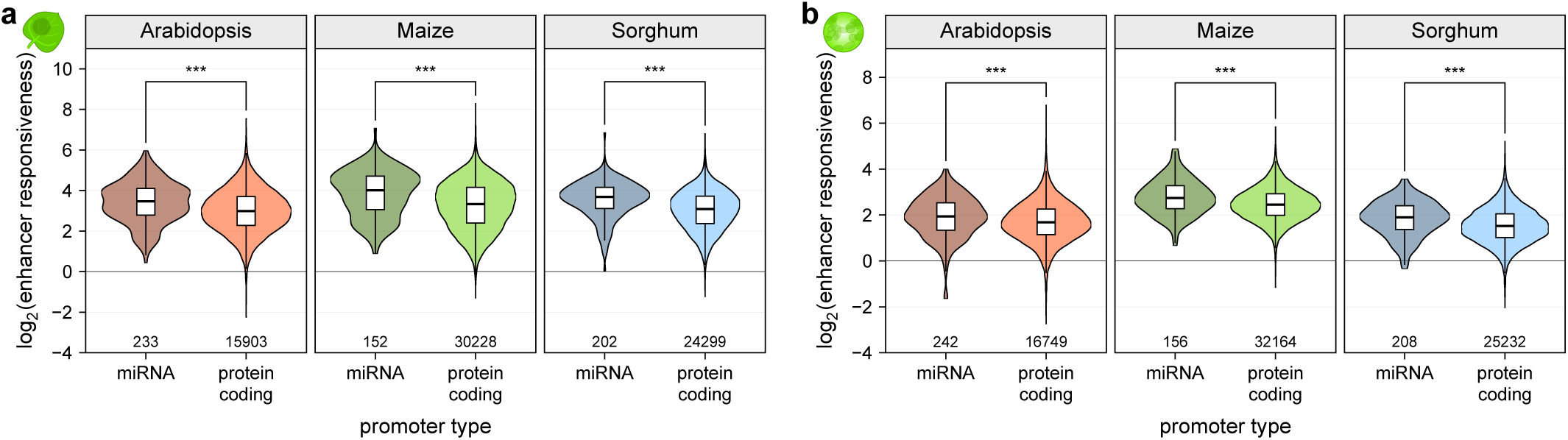
Promoters of miRNA genes are more responsive to the 35S enhancer than those of protein-coding genes. **a**,**b**, Violin plots (as defined in Figure 1) of enhancer responsiveness in tobacco leaves (**a**) or maize protoplasts (**b**). Promoters associated with miRNA or protein-coding genes are compared.

**Supplementary Fig. 13.**
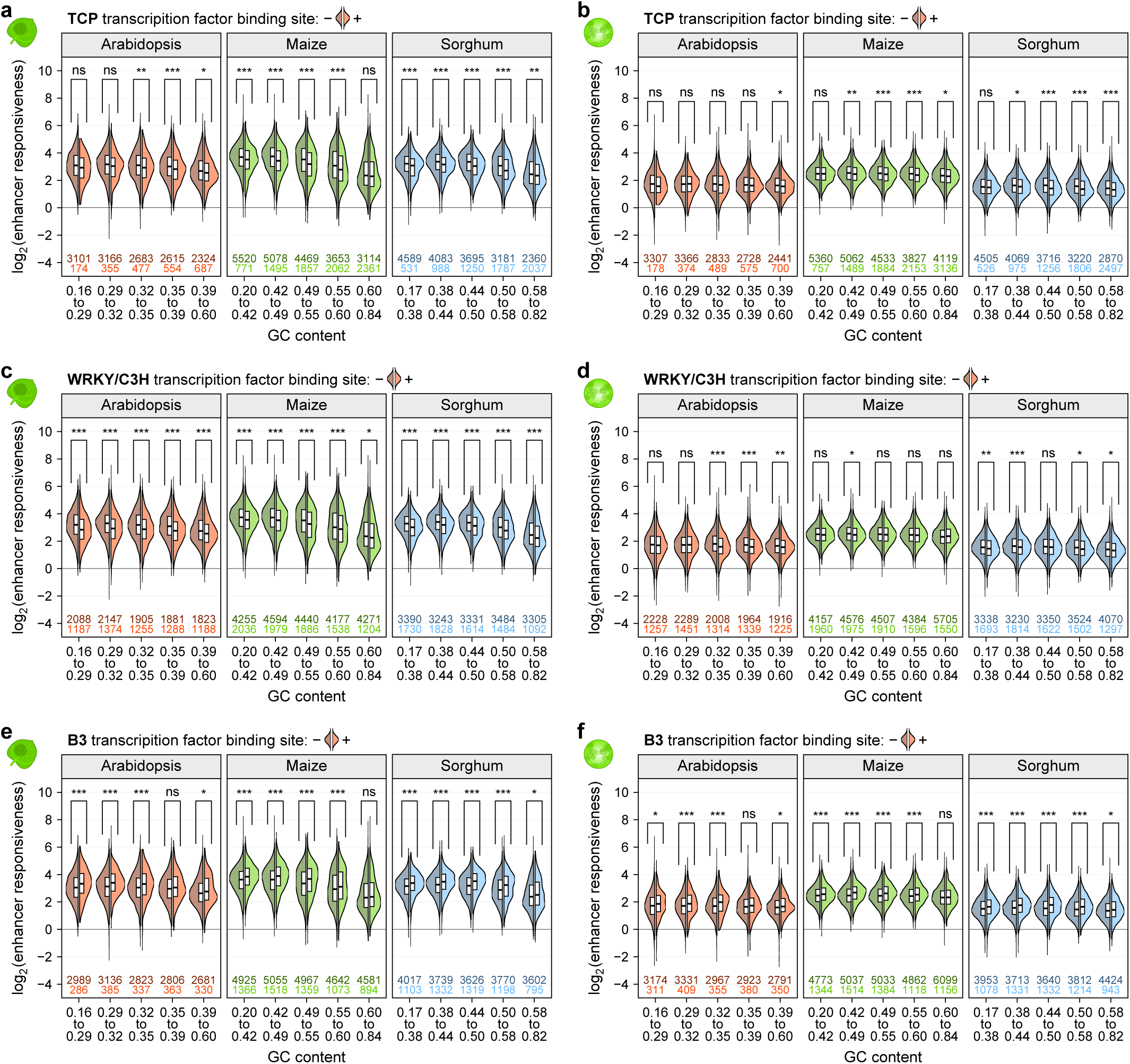
Promoter-proximal transcription factor binding sites influence enhancer responsiveness. **a**-**f**, Violin plots of enhancer responsiveness in tobacco leaves (**a,c,e**) or maize protoplasts (**b,d,f**). Promoters were grouped by GC content and split into promoters without (left half, darker color) or with (right half, lighter color) a TCP (**a,b**), WRKY (**c,d**), or B3 (**e,f**) transcription factor binding site. Violin plots are as defined in Figure 1, except only one half is shown.

**Supplementary Fig. 14.**
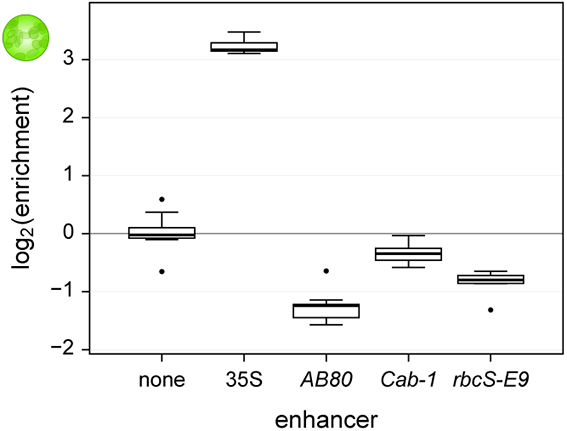
Light-responsive plant enhancers are not active in maize protoplasts. Constructs harboring no enhancer (none), a 35S enhancer, or one of three light-responsive plant enhancers (*AB80*, *Cab-1*, or *rbcS-E9*) upstream of the 35S minimal promoter were subjected to STARR-seq in maize protoplasts generated from dark-grown plants (Jores et al., 2020). Each boxplot (center line, median; box limits, upper and lower quartiles; whiskers, 1.5 × interquartile range; points, outliers) denotes the enrichment of all recovered mRNA barcodes over the DNA input. Only one experiment was performed.

**Supplementary Fig. 15.**
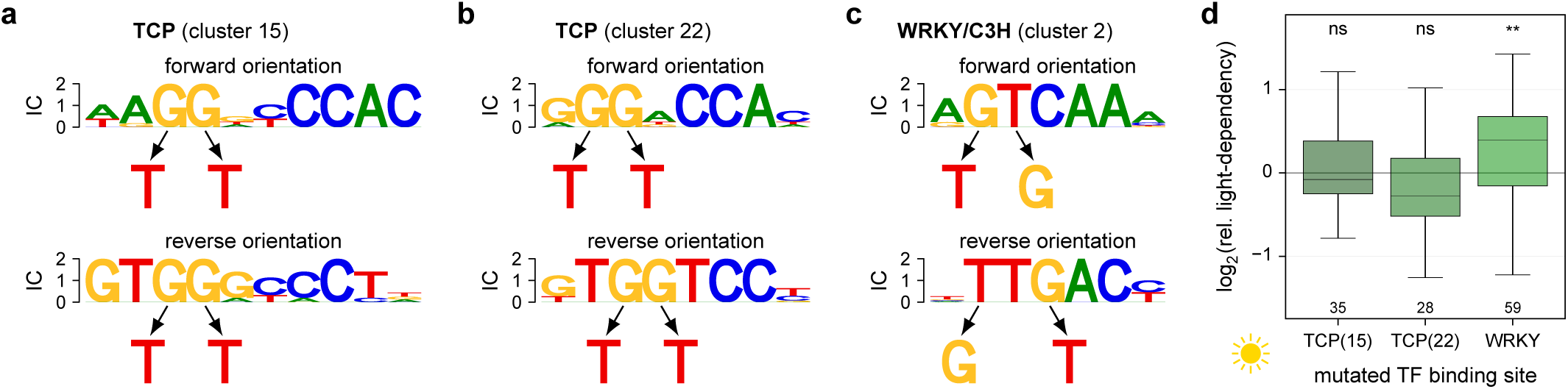
Mutations in transcription factor binding sites influence light-dependency. **a**-**c**, One or two T>G mutations were introduced in binding sites for TCP (**a,b**) or WRKY (**c**) transcription factors. Which bases were mutated depended on the orientation in which the binding site was present in the wild type promoter. **d**, Boxplots (as defined in Figure 3) for the relative light-dependency of promoters harboring mutations in the indicated transcription factor binding site as shown in (**a**-**c**). The corresponding wild type promoter was set to 0 (horizontal black line).

**Supplementary Fig. 16.**
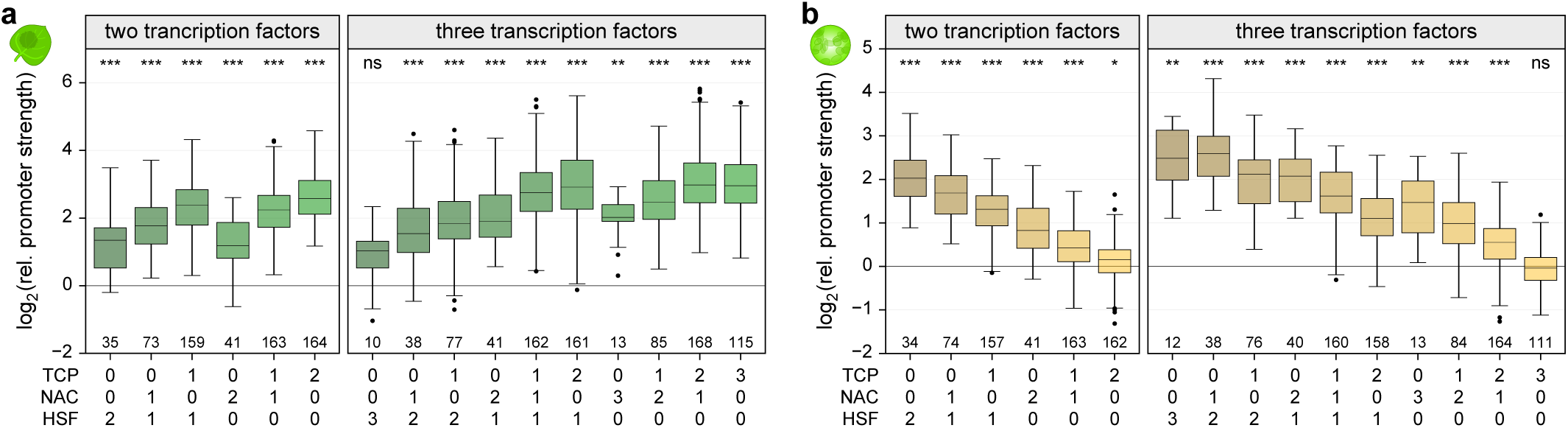
Transcription factor binding sites affect promoter strength additively. **a**,**b**, Boxplots (as defined in Figure 3) of promoter strength for libraries without enhancer in tobacco leaves (**a**) or maize protoplasts (**b**) for synthetic promoters with the indicated numbers of binding sites for TCP, NAC, and HSF transcription factors. The corresponding promoter without any transcription factor binding site was set to 0 (horizontal black line).

**Supplementary Fig. 17.**
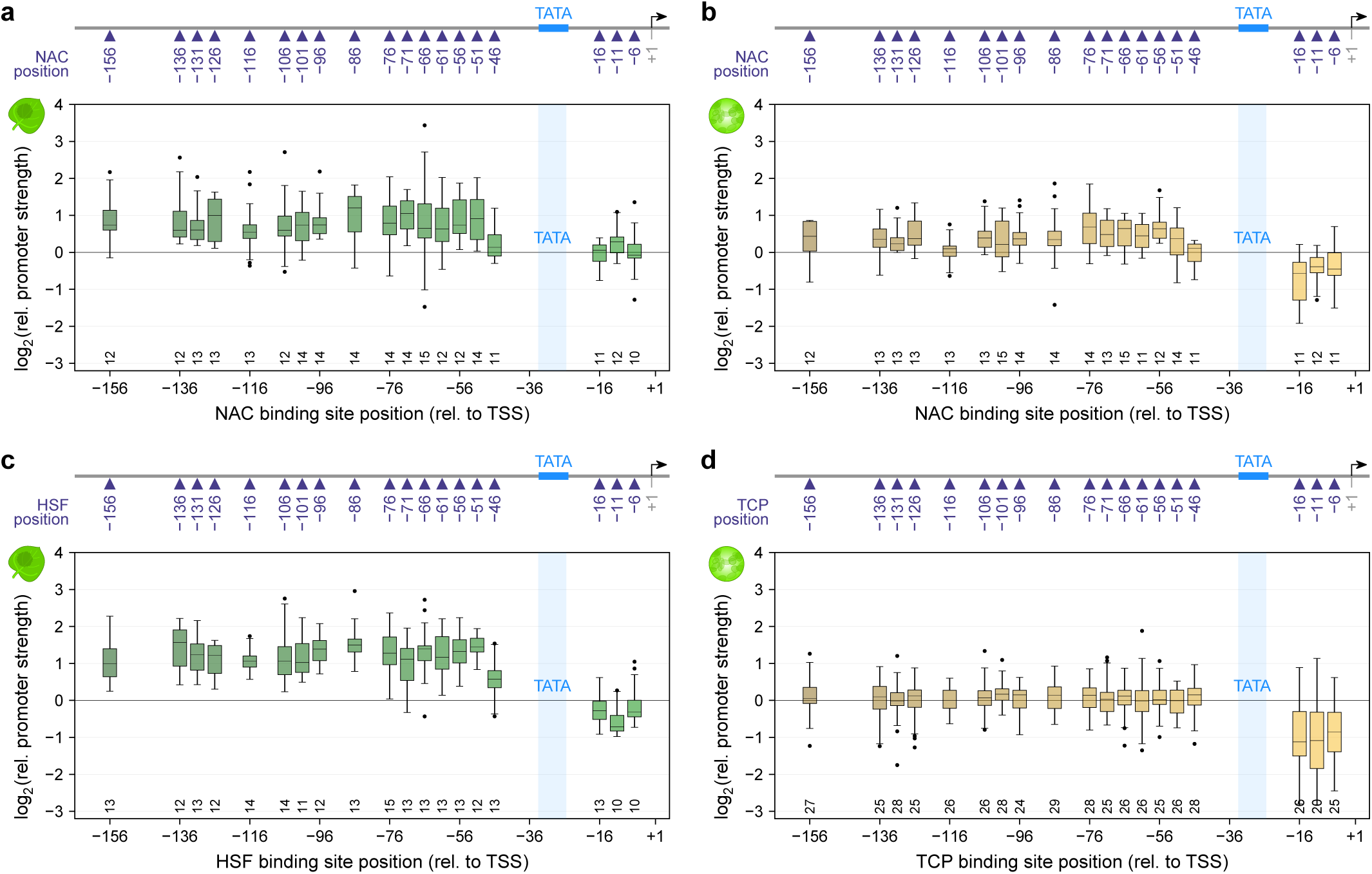
Transcription factor binding sites increase promoter strength only when present upstream of the TATA-box. **a**-**d**, A single NAC (**a,b**), HSF (**c**) or TCP (**d**) transcription factor binding site was inserted at the indicated position in the synthetic promoters containing a TATA-box. The strength of these promoters was measured in tobacco leaves (**a,c**) or maize protoplasts (**b,d**). Boxplots are as defined in Figure 3. The corresponding promoter without any transcription factor binding was set to 0 (horizontal black line).

**Supplementary Fig. 18.**
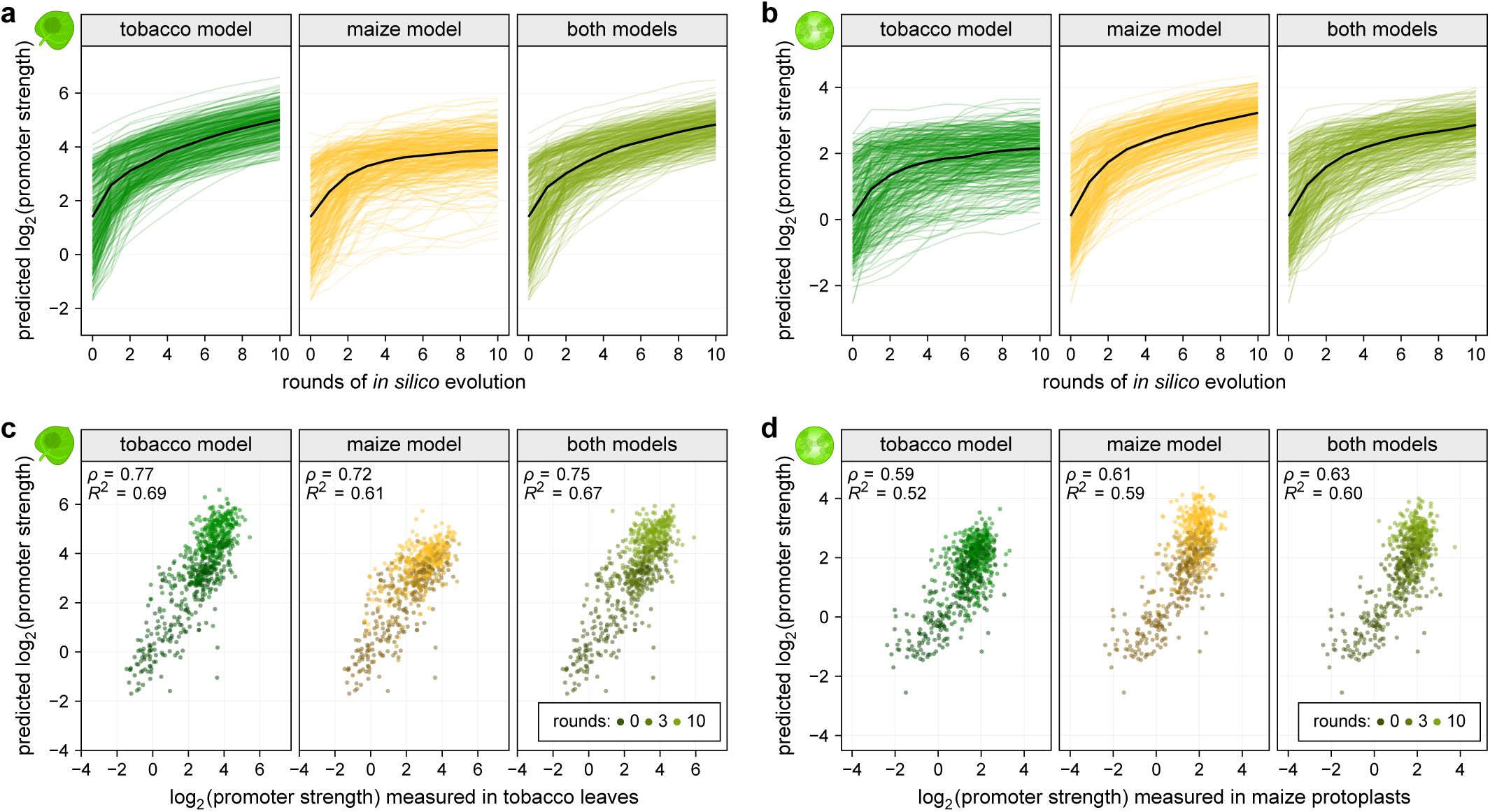
The *in silico* evolution is most effective in early rounds. **a**,**b**, 150 native and 160 synthetic promoters were subjected to 10 rounds of *in silico* evolution and the strength of the evolved promoters was predicted with the tobacco model (**a**) or the maize model (**b**). The black line represents the median promoter strength after each round. **c**,**d**, Correlation (Pearson’s *R*^2^ and Spearman’s *ρ*) between the predicted and experimentally-determined strength of promoters after 0, 3, or 10 rounds of *in silico* evolution. Promoter strengths measured in tobacco leaves were compared to predictions from the tobacco model (**c**) and the data from maize protoplasts was compared to the predictions from the maize model (**d**). The models used for the *in silico* evolution are indicated on each plot.

## Notes

### Competing Interest Statement

The authors have declared no competing interest.

## References

Andersson, R. and Sandelin, A. (2020). Determinants of enhancer and promoter activities of regulatory elements. Nat. Rev. Genet. 21: 71–87.

van Arensbergen, J., FitzPatrick, V.D., de Haas, M., Pagie, L., Sluimer, J., Bussemaker, H.J., and van Steensel, B. (2017). Genome-wide mapping of autonomous promoter activity in human cells. Nat. Biotechnol. 35: 145–153.

Arnold, C.D., Gerlach, D., Stelzer, C., Boryń, Ł.M., Rath, M., and Stark, A. (2013). Genome-Wide Quantitative Enhancer Activity Maps Identified by STARR-seq. Science 339: 1074–1077.

Arnold, C.D., Zabidi, M.A., Pagani, M., Rath, M., Schernhuber, K., Kazmar, T., and Stark, A. (2017). Genome-wide assessment of sequence-intrinsic enhancer responsiveness at single-base-pair resolution. Nat. Biotechnol. 35: 136–144.

Banerji, J., Olson, L., and Schaffner, W. (1983). A lymphocyte-specific cellular enhancer is located downstream of the joining region in immunoglobulin heavy chain genes. Cell 33: 729–740.

Banerji, J., Rusconi, S., and Schaffner, W. (1981). Expression of a β globin gene is enhanced by remote SV40 DNA sequences. Cell 27: 299–308.

Benfey, P.N., Ren, L., and Chua, N.H. (1990). Tissue-specific expression from CaMV 35S enhancer subdomains in early stages of plant development. EMBO J. 9: 1677–1684.

Bernard, V., Brunaud, V., and Lecharny, A. (2010). TC-motifs at the TATA-box expected position in plant genes: a novel class of motifs involved in the transcription regulation. BMC Genomics 11: 1–15.

Blake, M.C., Jambou, R.C., Swick, A.G., Kahn, J.W., and Azizkhan, J.C. (1990). Transcriptional initiation is controlled by upstream GC-box interactions in a TATAA-less promoter. Mol. Cell. Biol. 10: 6632–6641.

de Boer, C.G., Vaishnav, E.D., Sadeh, R., Abeyta, E.L., Friedman, N., and Regev, A. (2020). Deciphering eukaryotic gene-regulatory logic with 100 million random promoters. Nat. Biotechnol. 38: 56–65.

Bruce, W.B., Christensen, A.H., Klein, T., Fromm, M., and Quail, P.H. (1989). Photoregulation of a phytochrome gene promoter from oat transferred into rice by particle bombardment. Proc. Natl. Acad. Sci. 86: 9692–9696.

Burke, T.W. and Kadonaga, J.T. (1996). Drosophila TFIID binds to a conserved downstream basal promoter element that is present in many TATA-box-deficient promoters. Genes Dev. 10: 711–724.

Cheng, C.-Y., Krishnakumar, V., Chan, A.P., Thibaud-Nissen, F., Schobel, S., and Town, C.D. (2017). Araport11: a complete reannotation of the Arabidopsis thaliana reference genome. Plant J. 89: 789–804.

Cuperus, J.T., Groves, B., Kuchina, A., Rosenberg, A.B., Jojic, N., Fields, S., and Seelig, G. (2017). Deep learning of the regulatory grammar of yeast 5′ untranslated regions from 500,000 random sequences. Genome Res. 27: 2015–2024.

Deng, W. and Roberts, S.G.E. (2005). A core promoter element downstream of the TATA box that is recognized by TFIIB. Genes Dev. 19: 2418–2423.

Engler, C., Kandzia, R., and Marillonnet, S. (2008). A One Pot, One Step, Precision Cloning Method with High Throughput Capability. PLOS ONE 3: e3647.

Fang, R.X., Nagy, F., Sivasubramaniam, S., and Chua, N.H. (1989). Multiple cis regulatory elements for maximal expression of the cauliflower mosaic virus 35S promoter in transgenic plants. Plant Cell 1: 141–150.

Fornes, O. et al. (2020). JASPAR 2020: update of the open-access database of transcription factor binding profiles. Nucleic Acids Res. 48: D87–D92.

Gehrig, J., Reischl, M., Kalmár, É., Ferg, M., Hadzhiev, Y., Zaucker, A., Song, C., Schindler, S., Liebel, U., and Müller, F. (2009). Automated high-throughput mapping of promoter-enhancer interactions in zebrafish embryos. Nat. Methods 6: 911–916.

Grosschedl, R. and Birnstiel, M.L. (1980). Identification of regulatory sequences in the prelude sequences of an H2A histone gene by the study of specific deletion mutants in vivo. Proc. Natl. Acad. Sci. 77: 1432–1436.

Heerah, S., Katari, M., Penjor, R., Coruzzi, G., and Marshall-Colon, A. (2019). WRKY1 Mediates Transcriptional Regulation of Light and Nitrogen Signaling Pathways. Plant Physiol. 181: 1371–1388.

Hellens, R.P., Edwards, E.A., Leyland, N.R., Bean, S., and Mullineaux, P.M. (2000). pGreen: a versatile and flexible binary Ti vector for Agrobacterium-mediated plant transformation. Plant Mol. Biol. 42: 819–832.

Hong, C.K. and Cohen, B.A. (2021). Genomic environments scale the activities of diverse core promoters. bioRxiv: 2021.03.08.434469.

Ince, T.A. and Scotto, K.W. (1995). A Conserved Downstream Element Defines a New Class of RNA Polymerase II Promoters. J. Biol. Chem. 270: 30249–30252.

Jiao, Y. et al. (2017). Improved maize reference genome with single-molecule technologies. Nature 546: 524–527.

Jores, T., Tonnies, J., Dorrity, M.W., Cuperus, J.T., Fields, S., and Queitsch, C. (2020). Identification of Plant Enhancers and Their Constituent Elements by STARR-seq in Tobacco Leaves. Plant Cell 32: 2120–2131.

Kiran, K., Ansari, S.A., Srivastava, R., Lodhi, N., Chaturvedi, C.P., Sawant, S.V., and Tuli, R. (2006). The TATA-Box Sequence in the Basal Promoter Contributes to Determining Light-Dependent Gene Expression in Plants. Plant Physiol. 142: 364–376.

Klein, J.C., Agarwal, V., Inoue, F., Keith, A., Martin, B., Kircher, M., Ahituv, N., and Shendure, J. (2020). A systematic evaluation of the design and context dependencies of massively parallel reporter assays. Nat. Methods 17: 1083–1091.

Kotopka, B.J. and Smolke, C.D. (2020). Model-driven generation of artificial yeast promoters. Nat. Commun. 11: 2113.

Kumari, S. and Ware, D. (2013). Genome-Wide Computational Prediction and Analysis of Core Promoter Elements across Plant Monocots and Dicots. PLOS ONE 8: e79011.

Lagrange, T., Kapanidis, A.N., Tang, H., Reinberg, D., and Ebright, R.H. (1998). New core promoter element in RNA polymerase II-dependent transcription: sequence-specific DNA binding by transcription factor IIB. Genes Dev. 12: 34–44.

Lewis, B.A., Kim, T.-K., and Orkin, S.H. (2000). A downstream element in the human β globin promoter: Evidence of extended sequence-specific transcription factor IID contacts. Proc. Natl. Acad. Sci. 97: 7172–7177.

Lim, C.Y., Santoso, B., Boulay, T., Dong, E., Ohler, U., and Kadonaga, J.T. (2004). The MTE, a new core promoter element for transcription by RNA polymerase II. Genes Dev. 18: 1606–1617.

Liu, W. and Stewart, C.N. (2015). Plant synthetic biology. Trends Plant Sci. 20: 309–317.

Lomonossoff, G.P. and D’Aoust, M.-A. (2016). Plant-produced biopharmaceuticals: A case of technical developments driving clinical deployment. Science 353: 1237–1240.

Lubliner, S., Regev, I., Lotan-Pompan, M., Edelheit, S., Weinberger, A., and Segal, E. (2015). Core promoter sequence in yeast is a major determinant of expression level. Genome Res. 25: 1008–1017.

Madeira, F., Park, Y. mi, Lee, J., Buso, N., Gur, T., Madhusoodanan, N., Basutkar, P., Tivey, A.R.N., Potter, S.C., Finn, R.D., and Lopez, R. (2019). The EMBL-EBI search and sequence analysis tools APIs in 2019. Nucleic Acids Res. 47: W636–W641.

Masella, A.P., Bartram, A.K., Truszkowski, J.M., Brown, D.G., and Neufeld, J.D. (2012). PANDAseq: paired-end assembler for illumina sequences. BMC Bioinformatics 13: 1–7.

McCormick, R.F. et al. (2018). The Sorghum bicolor reference genome: improved assembly, gene annotations, a transcriptome atlas, and signatures of genome organization. Plant J. 93: 338–354.

Mejía-Guerra, M.K., Li, W., Galeano, N.F., Vidal, M., Gray, J., Doseff, A.I., and Grotewold, E. (2015). Core Promoter Plasticity Between Maize Tissues and Genotypes Contrasts with Predominance of Sharp Transcription Initiation Sites. Plant Cell 27: 3309–3320.

Mergner, J. et al. (2020). Mass-spectrometry-based draft of the Arabidopsis proteome. Nature 579: 409–414.

Molina, C. and Grotewold, E. (2005). Genome wide analysis of Arabidopsis core promoters. BMC Genomics 6: 25.

Morton, T., Petricka, J., Corcoran, D.L., Li, S., Winter, C.M., Carda, A., Benfey, P.N., Ohler, U., and Megraw, M. (2014). Paired-End Analysis of Transcription Start Sites in Arabidopsis Reveals Plant-Specific Promoter Signatures. Plant Cell 26: 2746–2760.

Onimaru, K., Nishimura, O., and Kuraku, S. (2020). Predicting gene regulatory regions with a convolutional neural network for processing double-strand genome sequence information. PLOS ONE 15: e0235748.

Parry, T.J., Theisen, J.W.M., Hsu, J.-Y., Wang, Y.-L., Corcoran, D.L., Eustice, M., Ohler, U., and Kadonaga, J.T. (2010). The TCT motif, a key component of an RNA polymerase II transcription system for the translational machinery. Genes Dev. 24: 2013–2018.

Patwardhan, R.P., Lee, C., Litvin, O., Young, D.L., Pe’er, D., and Shendure, J. (2009). High-resolution analysis of DNA regulatory elements by synthetic saturation mutagenesis. Nat. Biotechnol. 27: 1173–1175.

Raudvere, U., Kolberg, L., Kuzmin, I., Arak, T., Adler, P., Peterson, H., and Vilo, J. (2019). g:Profiler: a web server for functional enrichment analysis and conversions of gene lists (2019 update). Nucleic Acids Res. 47: W191–W198.

Rensink, W.A., Lee, Y., Liu, J., Iobst, S., Ouyang, S., and Buell, C.R. (2005). Comparative analyses of six solanaceous transcriptomes reveal a high degree of sequence conservation and species-specific transcripts. BMC Genomics 6: 1–14.

Ricci, W.A. et al. (2019). Widespread long-range cis-regulatory elements in the maize genome. Nat. Plants 5: 1237–1249.

Shahmuradov, I.A., Gammerman, A.J., Hancock, J.M., Bramley, P.M., and Solovyev, V.V. (2003). PlantProm: a database of plant promoter sequences. Nucleic Acids Res. 31: 114–117.

Sharon, E., Kalma, Y., Sharp, A., Raveh-Sadka, T., Levo, M., Zeevi, D., Keren, L., Yakhini, Z., Weinberger, A., and Segal, E. (2012). Inferring gene regulatory logic from high-throughput measurements of thousands of systematically designed promoters. Nat. Biotechnol. 30: 521–530.

Sheen, J. (1990). Metabolic repression of transcription in higher plants. Plant Cell 2: 1027–1038.

Singh, R., Ming, R., and Yu, Q. (2016). Comparative Analysis of GC Content Variations in Plant Genomes. Trop. Plant Biol. 9: 136–149.

Smale, S.T. and Baltimore, D. (1989). The “initiator” as a transcription control element. Cell 57: 103–113.

Smale, S.T. and Kadonaga, J.T. (2003). The RNA Polymerase II Core Promoter. Annu. Rev. Biochem. 72: 449–479.

Srivastava, A.K., Lu, Y., Zinta, G., Lang, Z., and Zhu, J.-K. (2018). UTR-Dependent Control of Gene Expression in Plants. Trends Plant Sci. 23: 248–259.

Srivastava, R., Rai, K.M., Srivastava, M., Kumar, V., Pandey, B., Singh, S.P., Bag, S.K., Singh, B.D., Tuli, R., and Sawant, S.V. (2014). Distinct Role of Core Promoter Architecture in Regulation of Light-Mediated Responses in Plant Genes. Mol. Plant 7: 626–641.

Tian, F., Yang, D.-C., Meng, Y.-Q., Jin, J., and Gao, G. (2020). PlantRegMap: charting functional regulatory maps in plants. Nucleic Acids Res. 48: D1104–D1113.

Tsai, F.T.F. and Sigler, P.B. (2000). Structural basis of preinitiation complex assembly on human Pol II promoters. EMBO J. 19: 25–36.

Walley, J.W., Sartor, R.C., Shen, Z., Schmitz, R.J., Wu, K.J., Urich, M.A., Nery, J.R., Smith, L.G., Schnable, J.C., Ecker, J.R., and Briggs, S.P. (2016). Integration of omic networks in a developmental atlas of maize. Science 353: 814–818.

Wang, B., Regulski, M., Tseng, E., Olson, A., Goodwin, S., McCombie, W.R., and Ware, D. (2018). A comparative transcriptional landscape of maize and sorghum obtained by single-molecule sequencing. Genome Res. 28: 921–932.

Wasylyk, B., Derbyshire, R., Guy, A., Molko, D., Roget, A., Téoule, R., and Chambon, P. (1980). Specific in vitro transcription of conalbumin gene is drastically decreased by single-point mutation in T-A-T-A box homology sequence. Proc. Natl. Acad. Sci. 77: 7024–7028.

Weingarten-Gabbay, S., Nir, R., Lubliner, S., Sharon, E., Kalma, Y., Weinberger, A., and Segal, E. (2019). Systematic interrogation of human promoters. Genome Res. 29: 171–183.

Yahraus, T., Chandra, S., Legendre, L., and Low, P.S. (1995). Evidence for a Mechanically Induced Oxidative Burst. Plant Physiol. 109: 1259–1266.

Yamamoto, Y.Y., Ichida, H., Abe, T., Suzuki, Y., Sugano, S., and Obokata, J. (2007). Differentiation of core promoter architecture between plants and mammals revealed by LDSS analysis. Nucleic Acids Res. 35: 6219–6226.

Yanai, I., Benjamin, H., Shmoish, M., Chalifa-Caspi, V., Shklar, M., Ophir, R., Bar-Even, A., Horn-Saban, S., Safran, M., Domany, E., Lancet, D., and Shmueli, O. (2005). Genome-wide midrange transcription profiles reveal expression level relationships in human tissue specification. Bioinformatics 21: 650–659.

Zhu, Q., Dabi, T., and Lamb, C. (1995). TATA box and initiator functions in the accurate transcription of a plant minimal promoter in vitro. Plant Cell 7: 1681–1689.

